# Myelin pathology is a key feature of X-linked Dystonia Parkinsonism

**DOI:** 10.1101/2025.10.07.680990

**Authors:** Priya Prakash, Kerry C Limberg, Weimin Zhang, Yu Zhao, Klaudia F Laborc, Anna O’Keeffe, Bryanna C Vilnaigre, Heather Appleby, Michael R O’Dea, Cara Fernandez-Cerado, Gierold Paul A Legarda, Michelle Sy, Edwin L Muñoz, Mark Angelo C Ang, Cid Czarina E Diesta, Justin Han, Ean Norenberg, Ellen B Penney, D Cristopher Bragg, Adam C Mar, Ran Brosh, Jef D Boeke, Shane A Liddelow

## Abstract

X-linked Dystonia-Parkinsonism (XDP) is a progressive, adult-onset neurodegenerative movement disorder that predominantly affects males of Filipino descent^1-3^. The disease is caused by the insertion of a SINE-VNTR-Alu subfamily F (SVA_F) retrotransposon within an intron of the TATA-box binding protein-associated factor 1 (*TAF1*) gene^4^. A major barrier to understanding the pathophysiology of XDP has been the lack of relevant animal models. Here, we introduce a novel conditional humanized XDP mouse model harboring a hybrid mouse-human *Taf1*/*TAF1* gene (hy*TAF1*) containing the pathogenic SVA_F insertion. We activated the hy*TAF1* in Nestin+ neural progenitor cells and found that the resulting XDP male mice recapitulate features of the human disease including severe motor impairment, striatal atrophy, and reactive gliosis. Transcriptomic, histological, and electron microscopy analysis revealed a dramatic reduction in oligodendrocyte lineage cells and widespread myelin disruption. Consistent with these findings, postmortem brain tissue from XDP patients revealed similar myelin pathology, including near-complete loss of myelin in parts of the medial prefrontal cortex. Together, these results identify oligodendrocyte dysfunction and myelin loss as previously unrecognized contributors to XDP pathogenesis, providing new mechanistic insight into this debilitating disorder.

## INTRODUCTION

X-linked Dystonia-Parkinsonism (XDP) is an adult-onset neurodegenerative movement disorder that exclusively affects individuals of Filipino ancestry, especially those from the island of Panay in the Philippines. It is caused by the insertion of a human-specific SINE-VNTR-*Alu* subfamily F (SVA_F) retrotransposon within intron 32 of the TATA-box binding protein-associated factor 1 (*TAF1*) gene^4^. *TAF1*, located on chromosome Xq13.1, consists of 38 highly conserved exons^5^ encoding a protein that forms the largest subunit of the multi-subunit transcription factor Transcription Factor IID. TAF1 plays a crucial role in recognizing the TATA box by interacting with TATA Binding Protein during transcription initiation by RNA polymerase II, making TAF1 indispensable for transcription regulation and cellular viability. The XDP-specific SVA_F element is a composite of an Alu-like sequence, a variable number of tandem repeats (VNTR) sequence, and a hexameric (CCCTCT)n repeat^6^. The length of this hexameric repeat varies among individuals and is inversely correlated with age of disease onset wherein longer repeats are associated with earlier symptom manifestation^4,7^. Clinically, patients present with progressive dystonia and late-onset parkinsonism, and mortality typically occurs within ten years of initial diagnosis^1,2^.

Although XDP primarily affects males, rare cases of symptomatic female carriers have been reported^8-10^. The disease has a prevalence of approximately 5.74 cases per 100,000 individuals in the Philippines^8^, with an average onset at 39.7 years^11^. Previous studies have identified the loss of striatal medium spiny neurons as a characteristic feature of the XDP brain^12,13^; however, the involvement of other brain regions isnot well defined. Another pathological feature of XDP is the presence of reactive glial cells, including astrocytes and microglia. Increased gliosis has been observed in XDP patient brain tissue, particularly in regions of the prefrontal cortex^14^, suggesting that other Central Nervous System (CNS) cell types may contribute to (and potentially exacerbate) neuronal loss and tissue pathology.

Our understanding of XDP pathophysiology is hindered by several key factors. First, since *TAF1* is expressed in all cells, the contribution of individual CNS cell types to disease pathology is not clearly defined. Identifying whether one or more cell types drive XDP pathology will enable us to focus on cell-type specific approaches to circumvent tissue pathology. Second, SVAs are hominid-specific and thus we have not been able to faithfully replicate disease phenotypes in a relevant non-primate in vivo model that preserves its fundamental genetic characteristics. Third, even though in vitro studies, including those using human induced pluripotent stem cells^15^, organoid models^6^, patient-derived cells^16^, and neuronal cell lines^17^ have provided insights into how the pathogenic SVA_F in *TAF1* leads to dysregulation of specific cell functions, these systems do not fully recapitulate the changes to the tissue environment during early and late developmental stages that are affected within specific regions of the disease tissue. A mouse model of XDP thus offers a powerful approach to investigate disease pathophysiology both during embryonic development and throughout adulthood, mirroring the developmental timeline and onset of symptoms seen in human patients. Moreover, such a model enables the study of cell-type specific contributions to disease pathology and enable us to probe the cell-type specific changes within specific CNS regions, offering critical insights into the spatial and temporal dynamics of XDP pathology.

Here, we introduce a newly engineered humanized model of XDP comprising a conditional mouse-human *TAF1* allele^18^. The hybrid *TAF1* (hy*TAF1*) comprises exons 1-24 of mouse *Taf1* and exons 25-38 of the human *TAF1* gene, including all intervening introns, and includes an XDP patient-derived SVA_F insertion, as well as an ancestral SVA_D (common to all humans), both of which are found in intron 32^18^. We generated a neural cell-specific mouse line wherein expression of the hybrid hy*TAF1* gene was induced specifically in Nestin+ progenitor and their daughter cells. We report that the male XDP mice carrying the Nestin-specific hy*TAF1* gene exhibit phenotypes that closely mirror those observed in human XDP patients. Specifically, these mice display markedly pronounced weight deficits as early as post-natal day 21 (P21) and complete mortality by P60. Furthermore, male XDP mice exhibit striatal tissue loss, lateral ventricle enlargement, and severe motor impairments such as gait abnormalities, impaired coordination and balance, and reduced grip strength. Notably, transcriptomic analysis of cortical and striatal tissue uncovered striking downregulation of oligodendrocyte-associated genes. These transcriptional changes were accompanied by pronounced myelin abnormalities in the cortex, striatum, and corpus callosum. Consistent with these findings, human postmortem brain tissue from XDP patients also exhibit myelin abnormalities in the striatum and nearly complete myelin loss in regions of the medial prefrontal cortex adjacent to the callosal sulcus. Together, our findings identify a previously unrecognized role for oligodendrocyte dysfunction and myelin dysregulation in XDP pathogenesis. Our work provides new mechanistic insight into XDP and establishes a valuable in vivo platform for investigating disease progression and potential therapeutic strategies.

## RESULTS

### Modeling XDP in mice by inserting the XDP-specific SVA_F into hyTAF1 in neural cells

We engineered a conditional partially humanized (hybrid) *Taf1* allele (Figure 1a). The hybrid gene (hy*TAF1*) comprises mouse *Taf1* exons 1-24 (with intervening introns) and human XDP patient-derived *TAF1* exons 25-38 (with intervening introns) and is only expressed following Cre-mediated recombination, a process we refer to as “conversion”. The 78 kb human *TAF1* genomic segment harbors both the XDP-specific SVA_F (hereafter SVA_F) and an ancestral SVA_D element (common to all humans) in intron 32 (Figure 1b). To mitigate potential cross-species incompatibility, nineteen divergent amino acids in the human TAF1 protein were substituted to match the mouse sequence, yielding a protein sequence identical to wild type mouse TAF1. Upon Cre-mediated recombination between two strategically located loxP sites, endogenous m*Taf1* exons 25-3’UTR (untranslated) region are excised and h*TAF1* exons 25-3’UTR are relocated immediately downstream of m*Taf1* exons 1-24, forming the hybrid gene.

**Figure 1.**
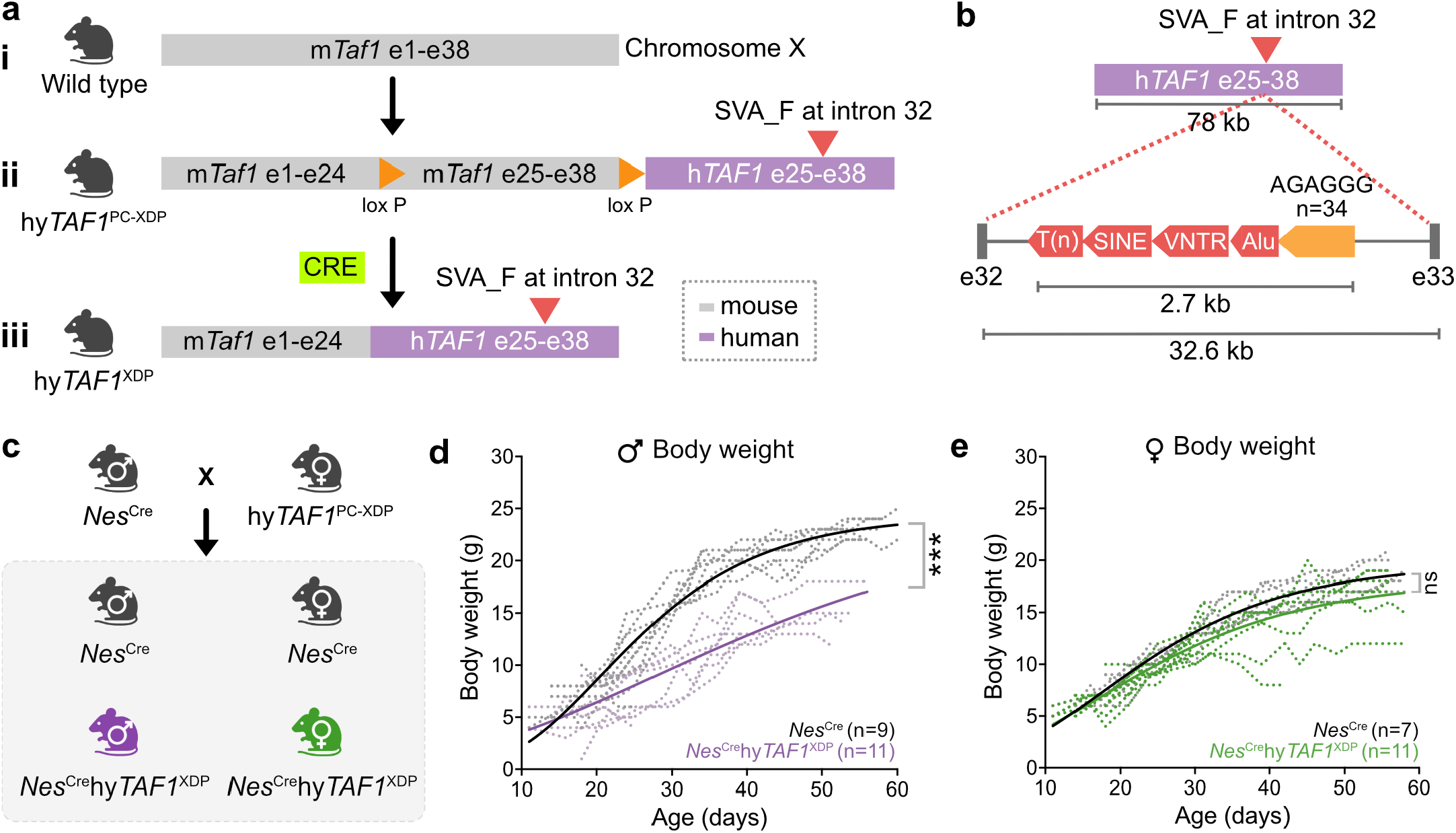
Genetic makeup of mouse/human hybrid hyTAF1 gene and generation of NesCrehyTAF1XDP mice. a. (i) Wild-type mouse on top with the full-length m*Taf1* spanning exons 1 to exons 38 (grey) on chromosome X. (ii) Introduction of an XDP patient-derived h*TAF1* segment spanning exons 25 to exons 38 (purple) downstream of m*Taf1*. An XDP-specific SVA_F is located in intron 32 of the h*TAF1* gene. Two loxP sites were inserted in the same orientation flanking the mouse exons 25-38 region. (iii) Cre-mediated excision of the m*Taf1* region results in a mouse-human hybrid *TAF1* gene. **b**. Structure of h*TAF1*. Composition of the disease-specific SVA_F element at intron 32 is shown. **c**. Crossing *Nes*^Cre^ homozygous males with hy*TAF1*^PC-XDP^ pre-converted XDP females results in *Nes*^Cre^ control male (black), *Nes*^Cre^hy*TAF1*^XDP^ male (purple), *Nes*^Cre^ control female (black), and *Nes*^Cre^hy*TAF1*^XDP^ carrier female (green) mice. **d**. Body weight curves of male *Nes*^Cre^ control (n = 9, black) and *Nes*^Cre^hy*TAF1*^XDP^ (n = 11, purple) mice. Dotted lines represent individual animals; bold lines indicate group means. Two-way ANOVA revealed significant main effects of genotype (p < 2 × 10^-16^) and age (p < 2 × 10^-16^) on body weight, as well as a significant genotype × age interaction (p = 0.00019), indicating that body weight changes over time differ between groups. Two-way ANOVA, *** p < 0.001. **e**. Body weight curves of female *Nes*^Cre^ control (n=7, black) and *Nes*^Cre^hy*TAF1*^XDP^ (n=11, green) mice. Dotted lines represent individual animals; bold lines indicate group means. Two-way ANOVA, trajectory of weight gain over time is non-significant (ns) between the two groups. Two-way ANOVA, p=0.0882.

To drive conversion of the conditional mouse/human hybrid hy*TAF1* in the CNS, we crossed female hy*TAF1*^PC-XDP^ mice with male Nestin^Cre^ (*Nes*^Cre^) mice. Nestin, an intermediate filament protein, is widely expressed by neural progenitor cells^19^ - enabling us to study the role of mutant *TAF1* in nearly all CNS cells. This cross generated four genotypes: *Nes*^Cre^ control males, *Nes*^Cre^hy*TAF1*^XDP^/Y males (representing human XDP patients), *Nes*^Cre^ control females, and *Nes*^Cre^hy*TAF1*^XDP^/+ females (representing human XDP carrier females) (Figure 1c). In *Nes*^Cre^ mice, Cre-mediated recombination occurs in Nestin+ progenitors and is maintained in their differentiated progeny^20^, allowing lineage-specific expression of hy*TAF1* in postmitotic neurons and other CNS cell types^20^.

Mendelian transmission of the *Taf1* allele was confirmed in offspring from these crosses (Supplementary Figure S1a). The observed distribution of genotypes was consistent with expected Mendelian ratios (∼25%), indicating that insertion of the SVA_F-containing h*TAF1* sequence into the mouse genome did not impair viability during embryonic development or result in preferential transmission of the allele (Supplementary Figure S1a). This confirms the stable inheritance and proper segregation of the modified *Taf1* allele in the presence and absence of Cre recombinase, enabling reliable generation of both experimental and control cohorts for downstream analysis.

We next evaluated the physical characteristics of the mice from all four genotypes. Compared to *Nes*^Cre^ control males *Nes*^Cre^hy*TAF1*^XDP^ males were visibly smaller in size (Supplementary Figure S1b), and some occasionally exhibited a hunched posture, ataxia, tremors (likely fatigue-induced), and tonic-clonic seizures. Longitudinal body weight measurements revealed a significant and progressive impairment in weight gain in *Nes*^Cre^hy*TAF1*^XDP^ males from postnatal day 21 (P21) to P60, relative to control males (Figure 1d, Supplementary Figure S1c). These mice also had a markedly reduced lifespan, with none surviving beyond P60^18^. Deaths were often sudden, with several cases likely resulting from seizure-related events and episodes resembling dystonic muscle spasms. In contrast, *Nes*^Cre^hy*TAF1*^XDP^ females did not show reduced body weights relative to female controls (Figure 1e, Supplementary Figure S1d) and survived beyond P60^18^.

In parallel with the hy*TAF1*^PC-XDP^ mice, we generated a control mouse line, hy*TAF1*^PC-ΔSVA_F^, which carries a similar hybrid allele composed of exons 1-24 from m*Taf1* and exons 25-38 from h^TAF1^ (Supplementary Figure S2a). However, unlike the disease model, the h*TAF1* segment in these mice lacks the XDP-specific SVA_F retrotransposon insertion in intron 32 (Supplementary Figure S2a). To activate the conditional hybrid allele in the CNS, we followed the same breeding scheme and generated *Nes*^Cre^hy*TAF1*^ΔSVA_F^ males. These mice express hy*TAF1* without the pathogenic SVA_F insertion specifically in Nestin+ neural lineages (Figure S2b). Unlike the *Nes*^Cre^hy*TAF1*^XDP^ males, which exhibit reduced body weight and early lethality, *Nes*^Cre^hy*TAF1*^ΔSVA_F^ males appeared phenotypically normal and maintained healthy body weights through postnatal development, including at P60, a time point by which 100% of the *Nes*^Cre^hy*TAF1*^XDP^ males had died (Figure S2c). Even at P67, *Nes*^Cre^hy*TAF1*^ΔSVA_F^ males displayed body weights comparable to those of age-matched *Nes*^Cre^ control littermates (Figure S2d).

To assess potential underlying tissue abnormalities, we collected brains from P90 *Nes*^Cre^ control and *Nes*^Cre^hy-*TAF1*^ΔSVA_F^ males and performed hematoxylin and eosin (H&E) staining to examine gross brain morphology. Both groups showed comparable tissue architecture, with no evidence of atrophy or histopathological abnormalities (Supplementary Figure S2e). These findings confirm that the insertion of the human *TAF1* genomic sequence alone, or the neuronal allele conversion early in development, do not cause the observed phenotypes in the disease model.

### Male XDP mice display severe motor impairments

We next conducted a comprehensive battery of behavioral assays to evaluate motor function in male and female *Nes*^Cre^hy*TAF1*^XDP^ mice across postnatal development from P14 to P35. Testing was organized into three developmental windows: week 3 (P14-21), week 4 (P21-28), and week 5 (P28-35) (Figure 2a).

**Figure 2.**
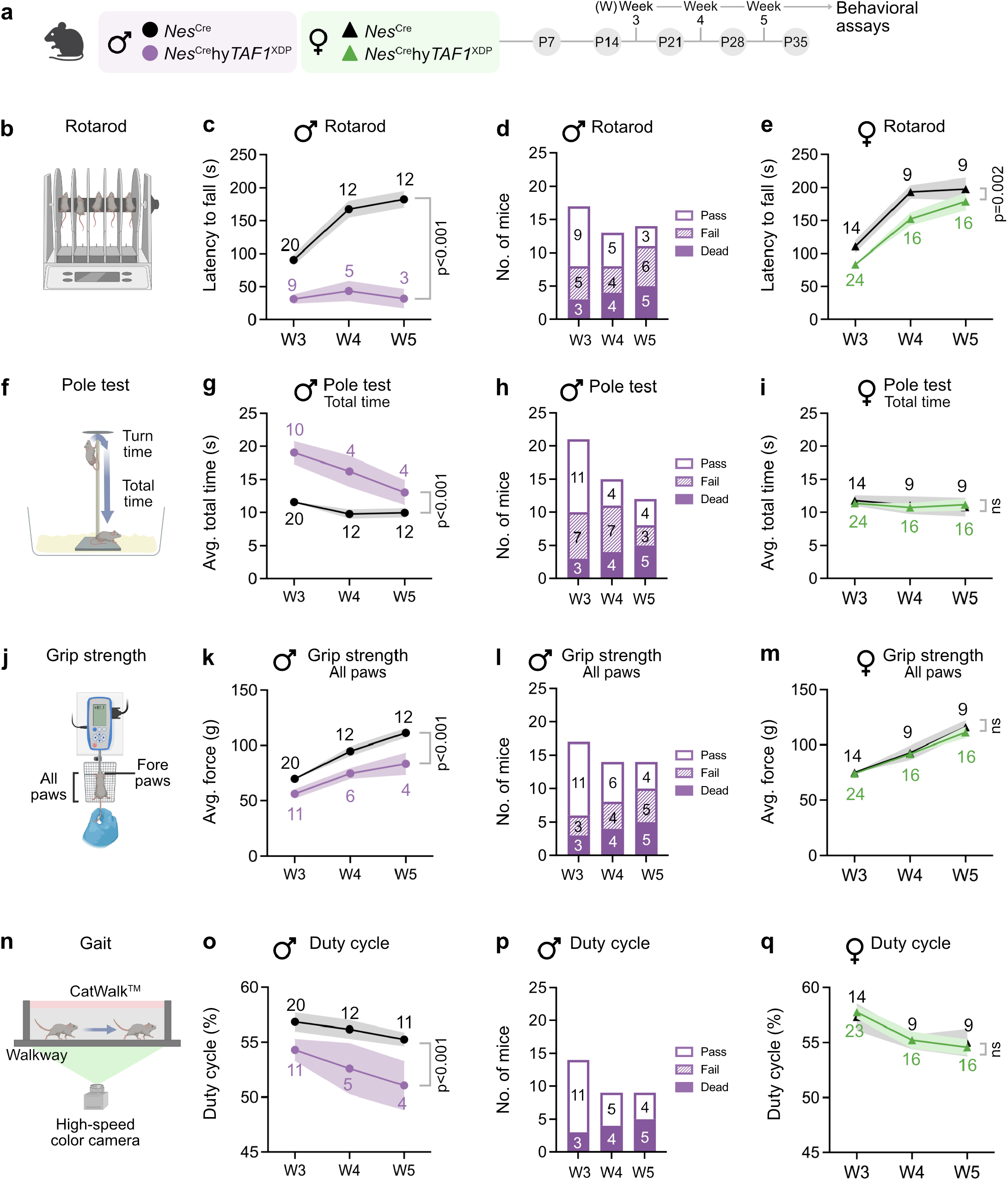
Motor function of *Nes*^Cre^hy*TAF1*^XDP^ male and female mice. **a**. Schematic illustrating the experimental timeline for assessing motor function in *Nes*^Cre^ and *Nes*^Cre^hy*TAF1*^XDP^ mice. ♂ (circles) and ♀ (triangles) mice underwent a battery of four behavioral assays designed to evaluate motor coordination and balance, grip strength, and gait. To monitor progression of motor function over time, these tests were performed at week (W) 3, 4, and 5. **b**. Rotarod test setup used to evaluate motor coordination and balance in mice. During the test, mice are placed on a rotating rod. The latency to fall, i.e., the time each mouse remains on the rod before losing balance and falling, is recorded. **c**. Latency to fall (s) of *Nes*^Cre^ and *Nes*^Cre^hy*TAF1*^XDP^ male mice. P < 0.001, linear mixed-effects model with age and timepoint as within-subject factors and genotype as a between-subject fixed effect. **d**. Number of *Nes*^Cre^hy*TAF1*^XDP^ mice that passed, failed, or died during rotarod test. Number of mice are displayed on the plot. **e**. Latency to fall (s) of *Nes*^Cre^ and *Nes*^Cre^hy-*TAF1*^XDP^ female mice. ns: non-significant. linear mixed-effects model with age and timepoint as within-subject factors and genotype as a between-subject fixed effect. **f**. Pole test. Mice are placed head-up at the top of a vertical pole, and two measures are recorded: time to orient downward (turn time) and total time to descend the pole (total time). **g**. Average total time (s) taken by *Nes*^Cre^ and *Nes*^Cre^hy*TAF1*^XDP^ male mice to descend the pole. p < 0.001, linear mixed-effects model with age and timepoint as within-subject factors and genotype as a between-subject fixed effect. **h**. Number of *Nes*^Cre^hy*TAF1*^XDP^ mice that passed, failed, or died during pole test. Number of mice are displayed on the plot. **i**. Average total time (s) taken by *Nes*^Cre^ and *Nes*^Cre^hy*TAF1*^XDP^ female mice to descend the pole. ns: non-significant, linear mixed-effects model with age and timepoint as within-subject factors and genotype as a between-subject fixed effect. **j**. Grip strength test. Mice grasp a bar connected to a force meter, and the peak force exerted is recorded. Grip strength is measured separately for forepaws alone and for all four paws. **k**. Average force (g) exerted by all paws by *Nes*^Cre^ and *Nes*^Cre^hy*TAF1*^XDP^ male mice. p < 0.001, linear mixed-effects model with age and timepoint as within-subject factors and genotype as a between-subject fixed effect. **l**. Number of *Nes*^Cre-^ hy*TAF1*^XDP^ mice that passed, failed, or died during assessment of grip strength. **m**. Average force (g) exerted by all paws by *Nes*^Cre^ and *Nes*^Cre^hy*TAF1*^XDP^ female mice. ns: non-significant, linear mixed-effects model with age and timepoint as within-subject factors and genotype as a between-subject fixed effect. **n**. Automated setup used to measure gait parameters in mice. As mice walk across an illuminated runway, a high-speed color camera positioned beneath captures images of their individual paw prints. **o**. Duty cycle (%) in male mice ((stand time)/(stand time+swing time)*100). Data were analyzed using linear mixed effects models with speed included as a covariate. Fixed effects were evaluated with Type III F-tests. Genotype had a significant effect on duty cycle (F(1, 21.40) = 28.72, p < 0.001), while neither time point (F(2, 14.65) = 0.21, p = 0.811) nor the genotype × time point interaction (F(2, 14.85) = 0.03, p = 0.968) were significant. The p-value shown in the plot represents the effect of genotype. **p**. Number of *Nes*^Cre^hy*TAF1*^XDP^ mice that passed, failed, or died during gait assessment using the CatWalk™ system. No mice failed the test. **q**. Duty cycle (%) in female mice ((stand time)/ (stand time+swing time)*100). Data were analyzed using linear mixed effects models with speed included as a covariate. Fixed effects were assessed using Type III F-tests. Genotype (F(1, 23.86) = 0.18, p = 0.673), time point (F(2, 22.38) = 0.40, p = 0.677), and the genotype × time point interaction (F(2, 22.99) = 1.16, p = 0.332) were not significant. “ns” on the plot indicates non-significant genotype differences.

To assess locomotor learning, coordination and balance, we used the rotarod test. Mice completed six trials each week, where they were placed on a rod that rotated under acceleration, gradually increasing from 4 to 40 rpm over 5 min. The time for each mouse to fall off, referred to as the “latency to fall”, was recorded (Figure 2b). Male *Nes*^Cre^hy*TAF1*^X-DP^ mice consistently exhibited poor performance compared to control *Nes*^Cre^ males, falling off the rod more quickly in every trial (Figure 2c, Supplementary Fiugure S3a-b). Test participation varied among mice; some were unable to perform and fell off the rod immediately, while others died during the test. These outcomes are summarized separately in a bar graph to illustrate the frequency of failures and mortality (Figure 2d). Female *Nes*^Cre^hy*TAF1*^XDP^ mice also exhibited slight motor deficits, showing reduced latency to fall compared to female *Nes*^Cre^ controls at corresponding time points (Figure 2e, Supplementary Figure S3c-d).

We used the pole test to assess limb strength, coordination and balance. Mice were placed head-up on top of a vertical pole, and two measures were recorded to assess motor coordination: the turn time, defined as the time taken for the mouse to orient itself downward, and the total time, which includes the full time taken to descend the pole to the base (Figure 2f). *Nes*^Cre^hy*TAF1*^XDP^ male mice exhibited significant impairments in this task too, taking longer to reorient and turn (Supplementary Figure S3g) and to descend the pole compared to *Nes*^Cre^ control males (Figure 2g, Supplementary Figure S3e). In addition, several *Nes*^Cre^hy*TAF1*^XDP^ males failed to complete the test, either by being unable to reorient themselves or by slipping and failing to descend properly, suggesting profound deficits in limb strength and motor coordination (Figure 2h). In contrast, female *Nes*^Cre^hy*TAF1*^XDP^ mice performed similarly to their *Nes*^Cre^ female counterparts, showing no significant impairment in either turn time or total descent time (Figure 2i, Supplementary Figure S3f,h).

Loss of grip strength, which affects hand function and manual dexterity, is a common symptom in patients with dystonia and Parkinson’s disease^21^. Mice underwent a grip strength test where we measured the maximal/peak force they exerted while grasping a metal grid with either their two fore paws or all four (fore and hind) paws, as they were pulled away (Figure 2j). *Nes*^Cre^hy*TAF1*^XDP^ male mice showed a significant reduction in average peak force (g) of both forepaws and fore- and hind-paws compared to *Nes*^Cre^ control males (Figure 2k, Supplementary Figure S3k,m). As with the other tests, a subset of *Nes*^Cre^hy*TAF1*^XDP^ mice failed the test by being unable to hold onto the grid, and a few animals died during the testing week (Figure 2l). In contrast, *Nes*^Cre^hy-*TAF1*^XDP^ female mice did not show any significant deficits in grip strength compared to female *Nes*^Cre^ controls (Figure 2m, Supplementary Figure S3l,n).

We assessed gait dynamics using the automated Cat-Walk™ system, where mice are allowed to walk freely across an illuminated glass walkway while a high-speed camera captures individual paw placements in real time (Figure 2n). This setup enables the extraction of numerous distinct gait parameters, including spatial and temporal characteristics of individual paws. One key parameter we analyzed was the duty cycle (%), which reflects the proportion of the step cycle during which a paw is in contact with the ground. We found that male *Nes*^Cre^hy*TAF1*^XDP^ mice exhibited significantly lower duty cycle (%) across all paws compared to *Nes*^Cre^ males (Figure 2o), after controlling for average speed. This indicates a fundamental difference in locomotor strategy of *Nes*^Cre^hy*TAF1*^XDP^ mice toward less supportive ground contact. The altered gait pattern may reflect compensation due to the smaller body size of *Nes*^Cre^hy*TAF1*^XDP^ mice and/or potential deficits in motor control, limb strength, balance or stability. All male mice successfully completed the Catwalk assay, with no test failures recorded (Figure 2p). In contrast to the males, female *Nes*^Cre^hy*TAF1*^XDP^ mice did not show any significant differences in duty cycle compared to female *Nes*^Cre^, suggesting preserved gait stability in females (Figure 2q).

Male *Nes*^Cre^hy*TAF1*^XDP^ mice also exhibited significant impairments in additional gait features. Specifically, they showed a reduction in paw position (mm) which reflects the placement of each step, and lower swing speed, indicating a slower movement of the paw during the swing phase of the step cycle (Supplementary Figure S4a,c). In contrast, female *Nes*^Cre^hy*TAF1*^XDP^ mice did not exhibit any significant differences in either paw position or swing speed when compared to their *Nes*^Cre^ female controls (Supplementary Figure S4b,d). We also examined additional gait parameters, including stride length, step cycle duration, cadence, and stance time. None of these metrics were significantly altered in either male or female *Nes*^Cre^hy*TAF1*^XDP^ mice compared to their respective controls (Supplementary Figure S4e-l). Together, these findings indicate that multiple aspects of gait are selectively affected in *Nes*^Cre^hy*TAF1*^XDP^ males, consistent with motor impairments reported in human XDP patients.

### XDP mouse brains exhibit signs of dystrophy and neuroinflammation

Given the more severe motor dysfunction observed in *Nes*^Cre^hy*TAF1*^XDP^ male mice at P21, we next examined their underlying tissue pathology and assessed markers of inflammation. These males often exhibited cranial swelling consistent with hydrocephaly. To evaluate gross tissue morphology, we performed H&E staining. At P21, *Nes*^Cre^ control males showed normal brain structure, whereas *Nes*^Cre^hy*TAF1*^XDP^ males displayed tissue loss, particularly in the striatum, along with enlarged lateral ventricles and degeneration of the dorsal striatal tissue (Figure 3a). Notably, there was variability in the severity of the tissue loss among individual mice: some exhibited relatively mild striatal tissue loss (Figure 3a, mouse 1), whereas others showed more extensive tissue loss, with one mouse demonstrating near-complete absence of the dorsal striatal tissue (Figure 3a, mouse 2,3).

**Figure 3.**
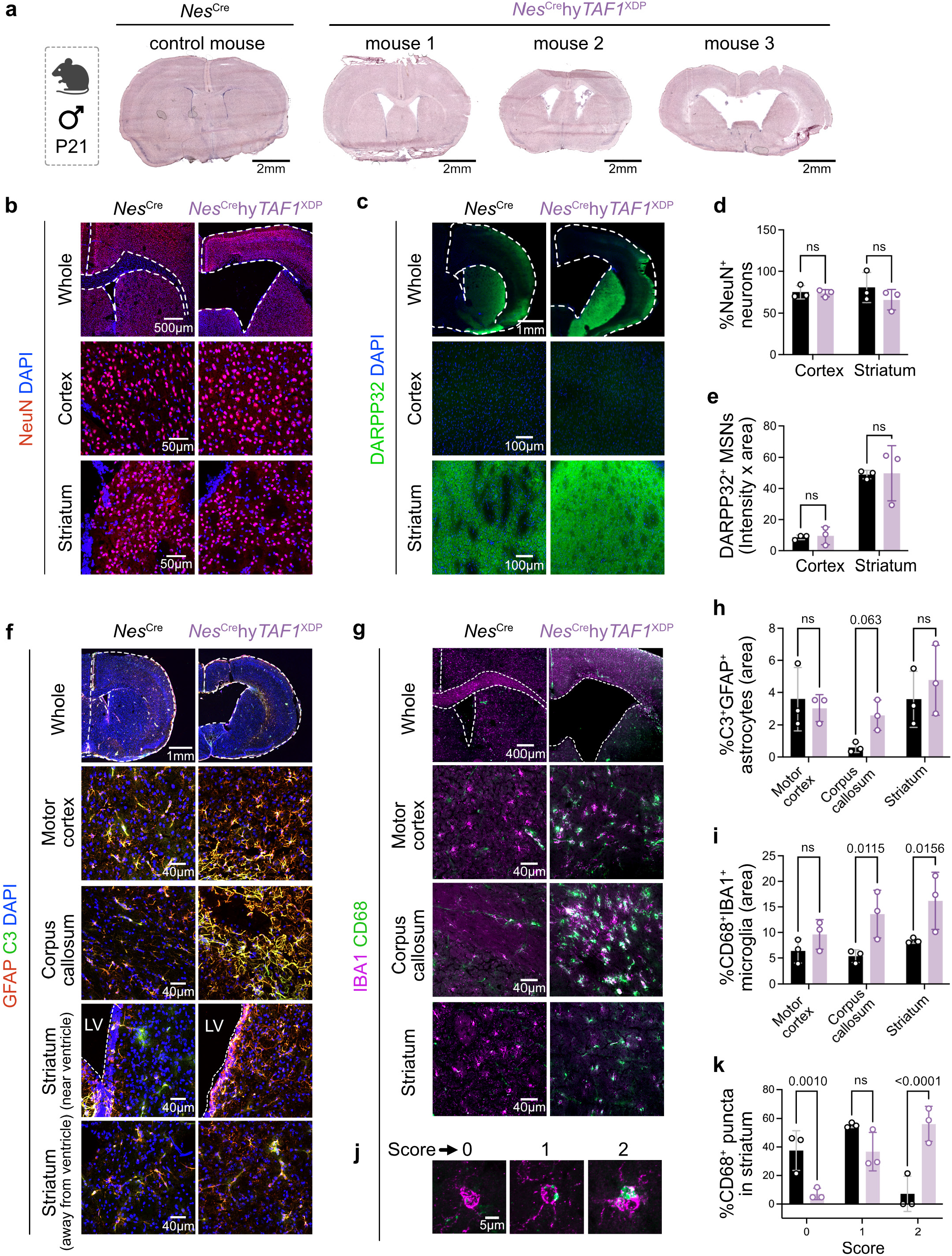
Immunohistochemical analysis reveals neuronal changes and glial reactivity in *Nes*^Cre^hy*TAF1*^XDP^ mouse brains. **a**. H&E staining of P21 male *Nes*^Cre^ control and three *Nes*^Cre^hy*TAF1*^XDP^ mouse brain tissues (left to right: less to more severity in pathology with enlarged lateral ventricles, and striatal tissue loss). Scale bar: 2 mm. **b**. Immunostaining for neurons using the pan-neuronal marker NeuN (red) in the cortex and striatum of P21 *Nes*^Cre^ and *Nes*^Cre^hy*TAF1*^XDP^ mice. Nuclei counterstained with DAPI (blue). **c**. Immunostaining for medium spiny neurons using DARPP-32 (green) in the cortex and striatum of P21 brains. Nuclei are counterstained with DAPI (blue). Scale bars: whole section = 1 mm; cortex and striatum panels = 100 µm. **d**. Quantification of neurons in the cortex and striatum as the percentage of NeuN+ cells relative to total DAPI+ nuclei (NeuN+/DAPI+ × 100). ns: non-significant, two-way ANOVA. **e**. Quantification of medium spiny neurons in cortex and striatum based on DARPP-32 signal, calculated as fluorescence intensity multiplied by stained area (integrated density). ns: non-significant, two-way ANOVA. **f**. Immunostaining of the motor cortex, corpus callosum, and striatum for astrocyte reactivity. Astrocytes labeled with GFAP (marker of astrocytes), C3 (marker of reactive astrocytes), and DAPI (nuclei). Scale bars: whole brain = 1 mm; region panels = 100 µm. **g**. Immunostaining of the motor cortex, corpus callosum, and striatum for microglial reactivity. Microglia labeled with IBA1 (pan-microglial marker) and CD68 (lysosomal marker of reactive microglia). Scale bars: whole section = 400 µm; region panels = 40 µm. **h**. Quantification of reactive astrocytes as the percentage of GFAP+C3+ double-positive cells over total GFAP+ cells in each brain region (motor cortex, corpus callosum, striatum). **i**. Quantification of reactive microglia as the percentage of IBA1+ area that co-localizes with CD68 signal: (CD68+IBA1+ overlap area/IBA1+ area) × 100. n = 3 mice per genotype. ns: non-significant, p values on the plot, two-way ANOVA. **j**. Representative images of CD68 puncta in IBA1+ microglia, showing scoring criteria: Score 0 = no puncta per cell; Score 1 = 1–3 puncta per cell; Score 2 = >2 puncta and/or enlarged puncta per cell. **k**. Quantification of CD68 puncta scores in IBA1+ microglia within the striatum. Data represent mean scores per group across multiple microglia and mice. n = 3 mice per genotype. ns: non-significant, p values on the plot, two-way ANOVA.

To determine how striatal tissue loss relates to changes in neuronal numbers within the remaining tissue, we performed immunohistochemical analyses using neuronal markers. First, we used NeuN as a pan-neuronal marker to assess changes in all neurons in both vulnerable (striatal) and non-vulnerable (cortical) regions. In the cortex and striatum, there was no difference in the number of NeuN+ cells between *Nes*^Cre^ and *Nes*^Cre^hy*TAF1*^XDP^ male mice (Figure 3b,d). Because portions of the striatal tissue were already lost at this time point (P21), our analysis focused on the striatal areas that remained intact. Next, we examined dopamine and cAMP-regulated phosphoprotein (DARPP-32), which is uniquely expressed by medium spiny neurons. As expected, DARPP-32 was absent from cortical regions and present within striatal tissue (Figure 3c). In the remaining striatal tissue of *Nes*^Cre^hy*TAF1*^XDP^ mice, we did not observe a significant reduction in DARPP-32 intensity compared to controls (Figure 3c-d). Together, these data suggests that there is no significant change occurring to the neuronal cell populations in the cortex as well as in the remainder of the striatal tissue at the examined timepoint in the *Nes*^Cre^hy*TAF1*^XDP^ male mice.

Because neuroinflammation is a hallmark of many CNS pathologies, we next examined changes in classical cellular markers of neuroinflammation. Reactive astrocytes are typically found near sites of neuronal death, and a subtype of reactive astrocytes characterized by increased C3 expression has been identified in several different pathological conditions. Given the level of neurodegeneration in these brain regions, we assessed the presence of C3+GFAP+ astrocytes in vulnerable striatal tissue, less vulnerable cortical areas, and white matter-rich regions such as the corpus callosum (Figure 3f). In the motor cortex, there was no significant difference in the number of C3+GFAP+ astrocytes between *Nes*^Cre^hy*TAF1*^XDP^ and *Nes*^Cre^ male mice (Figure 3h). Although we observed numerous C3+GFAP+ astrocytes immediately surrounding the lateral ventricles in the corpus callosum and striatum of *Nes*^Cre^hy*TAF1*^XDP^ mice, these differences were not statistically significant (Figure 3h).

Additionally, we evaluated microglial reactivity within these brain regions (Figure 3g). Reactive microglia are commonly observed in atrophied tissue in various neurodegenerative and neurological disorders^22^. They are characterized by increased expression of the lysosomal and phagocytic marker CD68. We did not observe a significant increase in the percentage of CD6+IBA1+ cells in the motor cortex of *Nes*^Cre^hy*TAF1*^XDP^ male mice compared to *Nes*^Cre^ controls. However, we did find a significant increase in CD68+IBA1+ cells specifically within the striatum and the white matter-rich corpus callosum (Figure 3i), suggesting heightened microglial reactivity. To further assess microglial reactivity, we analyzed the number and size of CD68 puncta within individual IBA1+ microglia. We assigned a score of 0 for no puncta, 1 for 1-3 puncta per microglial cell, and 2 for more than 3 puncta or large, diffuse puncta throughout the cell body (Figure 3j). In the striatum, corpus callosum, and motor cortex, IBA1+ microglia from *Nes*^Cre^hy*TAF1*^XDP^ mice exhibited significantly more, larger and more diffuse CD68+ puncta per microglia (score 3) (Figure 3k, Supplementary Figure S5a-c), consistent with increased microglial reactivity in these regions.

Taken together, these results demonstrate that at P21, *Nes*^Cre^hy*TAF1*^XDP^ male mice exhibit pronounced striatal atrophy accompanied by increased markers of glial reactivity and neuroinflammation particularly within white matter regions.

### Oligodendrocyte genes are significantly downregulated in the XDP mouse brain

We next used RNA sequencing to better characterize the response of CNS cells to XDP pathology. To investigate these molecular changes in male *Nes*^Cre^hy*TAF1*^XDP^ mice, we dissected the cortex and striatum from P21 *Nes*^Cre^hy*TAF1*^XDP^ and *Nes*^Cre^ control mice, isolated RNA, and performed bulk RNA sequencing (Figure 4a). Principal component analysis (PCA) revealed a clear separation between *Nes*^Cre^hy*TAF1*^XDP^ and *Nes*^Cre^ samples in both cortical and striatal datasets (Figure 4b-c), indicating distinct transcriptomic profiles between genotypes.

**Figure 4.**
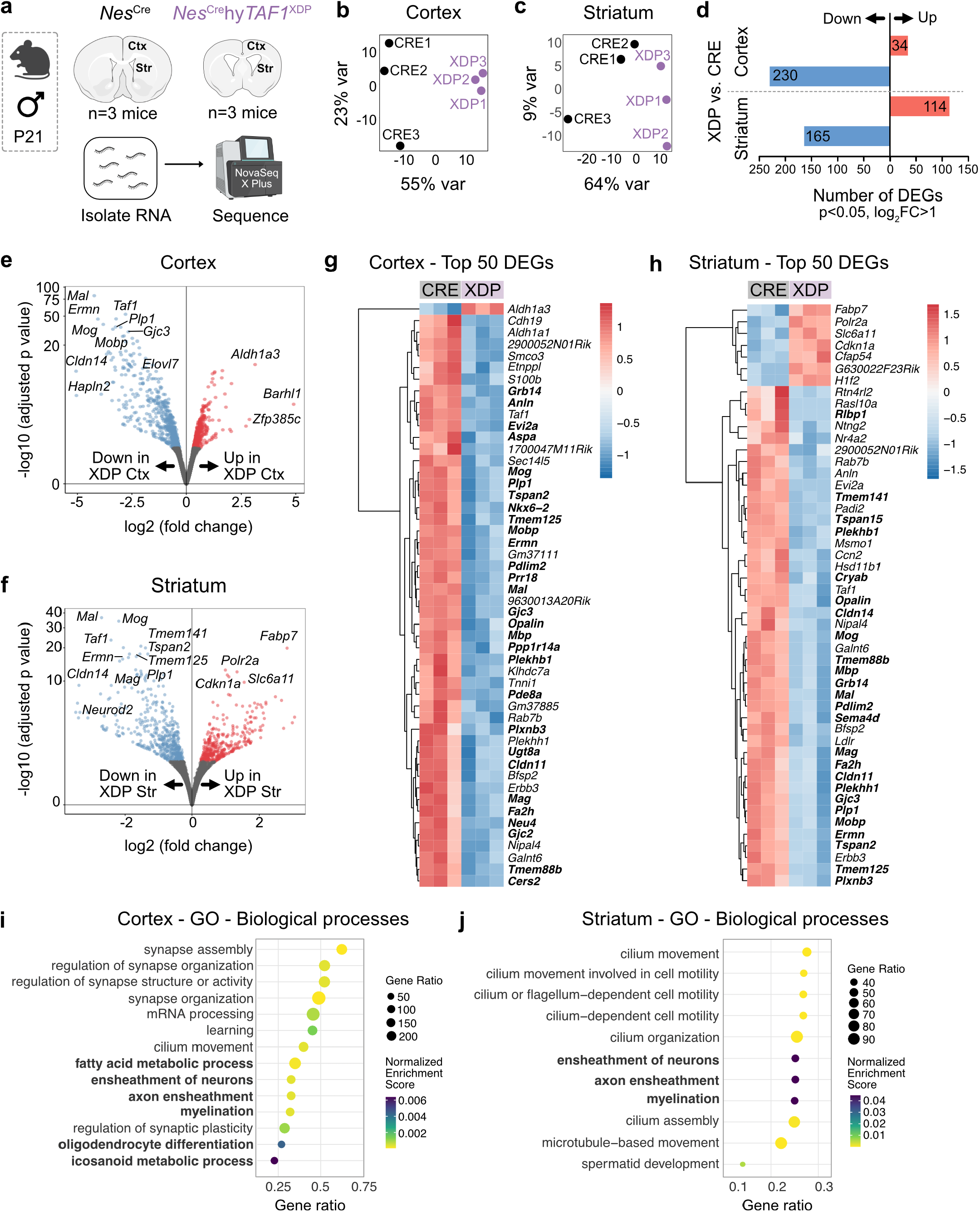
Transcriptomic analysis of *Nes*^Cre^ and *Nes*^Cre^hy*TAF1*^XDP^ cortical and striatal tissue by bulk RNA sequencing. **a**. RNA sequencing workflow. Cortical and striatal tissue was micro-dissected from P21 male *Nes*^Cre^ and *Nes*^Cre^hy*TAF1*^XDP^ mice (n = 3 per genotype). Total RNA was extracted and subjected to bulk RNA sequencing. **b**. Principal component analysis (PCA) of cortical transcriptomes for *Nes*^Cre^ (CRE) and *Nes*^Cre^hy*TAF1*^XDP^ (XDP) samples. **c**. PCA of striatal transcriptomes for *Nes*^Cre^ (CRE) and *Nes*^Cre^hy*TAF1*^XDP^ (XDP) samples. **d**. Number of differentially expressed genes (DEGs) in the cortex and striatum (adjusted p < 0.05, log_2_ fold change > 1). Red and blue bars represent up- and down-regulated genes, respectively. **e**. Volcano plot showing the distribution of log_2_ fold changes and −log_10_ adjusted p values for DEGs in the cortex. Selected genes, including those enriched in oligodendrocytes (*Mal, Ermn, Plp1, Mog, Mobp*, etc.), are highlighted. **f**. Volcano plot showing the distribution of log_2_ fold changes and −log_10_ adjusted p values for DEGs in the striatum, with selected oligodendrocyte-associated downregulated genes labeled. **g**. Heatmap of the top 50 DEGs in the cortex ranked by adjusted p-value. Expression values are z-scored per gene across all samples. Oligodendrocyte genes are bolded. **h**. Heatmap of the top 50 DEGs in the striatum ranked by adjusted p-value. Expression values are z-scored per gene across all samples. Oligodendrocyte genes are bolded. **i**. Gene Set Enrichment Analysis (GSEA) of cortical DEGs (FDR q < 0.25). Myelin- and oligodendrocyte-associated gene sets are highlighted in bold. **j**. GSEA of striatal DEGs (FDR q < 0.25).

Within the cortex, we identified 34 significantly upregulated and 230 significantly downregulated genes in *Nes*^Cre-^ hy*TAF1*^XDP^ mice compared to *Nes*^Cre^ controls (adjusted p < 0.05, log_2_ fold change > 1) (Figure 4d-e). Among the down-regulated genes, *Cldn14, Hapln2, Mal*, and *Ermn* showed the most dramatic decreases (Figure 4e-g). A majority of the significantly downregulated genes, including *Mag, Plp1, Mog*, and *Gjc2*, are highly enriched in oligodendrocytes (OLs), the myelinating glial cells of the CNS. Strikingly, several OL-enriched genes, including *Mag, Mal, Plp1, Mog*, and *Ermn*, were also significantly downregulated in the *Nes*^Cre-^ hy*TAF1*^XDP^ striatum (Figure 4d,f,h). *Taf1* itself was also significantly downregulated in both the cortex and striatum of *Nes*^Cre^hy*TAF1*^XDP^ mice (Figure 4e-f). While genes associated with mature OLs were significantly downregulated in both *Nes*^Cre^hy*TAF1*^XDP^ cortex and striatum, genes associated with OL precursor cells (OPCs) and pre-myelinating OLs were not significantly altered. These included *Ng2, Olig1/2, Gpr17/37/56, Sox2/5/6/8/10, Myrf*, among others. These results suggest a deficit in maturation and/or differentiation of OPCs into mature myelinating OLs.

To identify biological pathways associated with gene expression changes in the cortex and striatum of XDP mice compared to Cre controls, we performed Gene Set Enrichment Analysis (GSEA) using ranked log_2_ fold changes and Gene Ontology Biological Process annotations. GSEA revealed significant enrichment of terms related to OL function, including ensheathment of neurons, oligodendrocyte differentiation, and myelination (Figure 4i,j). These findings further support the notion that dysregulation of OL-associated pathways may contribute to regional tissue vulnerability in XDP mice.

We also performed bulk RNA sequencing on cortex and striatum of *Nes*^Cre^hy*TAF1*^ΔSVA_F^ and corresponding *Nes*^Cre^ control males (Supplementary Figure S6a). In contrast to the marked transcriptomic differences observed between *Nes*^Cre-^ hy*TAF1*^XDP^ and *Nes*^Cre^ control males, PCA of both cortical and striatal datasets revealed no clear separation between *Nes*^Cre^hy*TAF1*^ΔSVA_F^ and control samples (Supplementary Figure S6b-c). In the cortex, only 33 genes were upregulated and 27 downregulated in *Nes*^Cre^hy*TAF1*^ΔSVA_F^ mice compared to *Nes*^Cre^ controls (adjusted p < 0.05, log^2^ fold change > 1) (Supplementary Figure S6d-e,g), indicating minimal transcriptional disruption. In the striatum, the effect was even more limited, with no upregulated genes and only a single downregulated gene, *Taf1*, detected between *Nes*^Cre^hy*TAF1*^ΔSVA_F^ and *Nes*^Cre^ groups under the same statistical threshold (Supplementary Figure S6d,f,h). These findings suggest that deletion of the XDP-specific SVA_F element, while retaining the hybrid mouse/human TAF1 (hy*TAF1*), does not induce widespread transcriptional changes associated with XDP at this time-point.

### Aberrant myelination is a hallmark of the XDP mouse brain

To validate the transcriptomic findings implicating OLs, we next analyzed the *Nes*^Cre^hy*TAF1*^XDP^ and *Nes*^Cre^ control brain tissue to assess myelination and changes in oligodendrocyte-lineage cells using immunohistochemistry. Compared to *Nes*^Cre^ control males, *Nes*^Cre^hy*TAF1*^XDP^ male mice exhibited disrupted patterning of myelin oligodendrocyte glycoprotein (MOG) and myelin basic protein (MBP), membrane proteins expressed on the surface of oligodendrocytes and the outer-most layer of myelin sheaths. In the cortex and corpus callosum, the typical fibrous-like distribution of MOG and MBP were markedly altered in XDP mice (Figure 5a, Supplementary Figure S7a). Additionally, MOG and MBP staining in the striatum, which are normally seen as a distinct ‘halo’ at the borders of the striosomes, were also disrupted. In XDP mice, MOG and MBP appeared as disorganized, patchy pockets within the striosomes rather than being restricted to their borders, indicating abnormal myelination in these vulnerable regions (Figure 5a, Supplementary Figure S7a-b).

**Figure 5.**
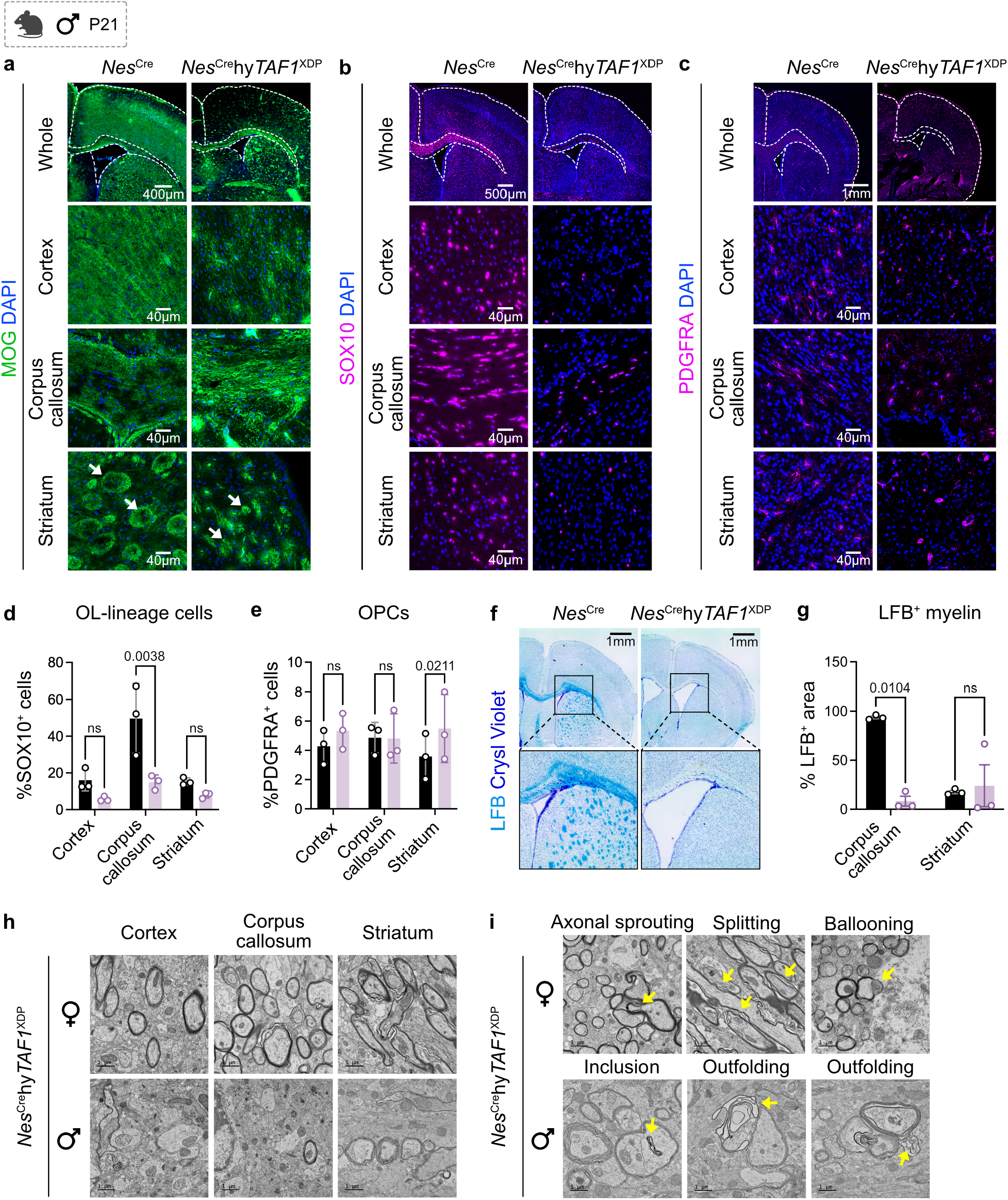
Myelin pathology and oligodendrocyte-lineage abnormalities in *Nes*^Cre^hy*TAF1*^XDP^ mouse brains revealed by immunohistochemistry and electron microscopy. **a**. Immunostaining of the motor cortex, corpus callosum, and striatum for Myelin Oligodendrocyte Glycoprotein (MOG, green), showing disrupted and patchy myelination in *Nes*^Cre^hy*TAF1*^XDP^ brains compared to *Nes*^Cre^ controls. Scale bars: whole section = 400 µm; region panels = 40 µm. **b**. Immunostaining for oligodendrocyte lineage cells using SOX10 (magenta) and DAPI (blue) in motor cortex, corpus callosum, and striatum. Scale bars: whole = 500 µm; region = 40 µm. **c**. Immunostaining for oligodendrocyte precursor cells (OPCs) using PDGFRA (magenta) and DAPI (blue) in the same regions. Scale bars: whole = 1 mm; region = 40 µm. **d**. Quantification of SOX10+ cells (% of DAPI+) in cortex, corpus callosum, and striatum in *Nes*^Cre^ and *Nes*^Cre^hy*TAF1*^XDP^ brains. Two-way repeated measures ANOVA with Šídák’s multiple comparisons test; p = 0.0038 (**), cortex p > 0.05 (ns), striatum p > 0.05 (ns). Data shown as mean +/− SEM; n = 3 mice/genotype. **e**. Quantification of PDGFRA+ cells (% of DAPI+) across brain regions. Two-way repeated measures ANOVA with Šídák’s multiple comparisons test; striatum p = 0.0211 (*), cortex and corpus callosum p > 0.05 (ns). Data shown as mean +/− SEM; n = 3 mice/genotype. **f**. Luxol Fast Blue (LFB) staining of myelin lipids in large brain sections. *Nes*^Cre^hy*TAF1*^XDP^ brains show decreased LFB staining in the corpus callosum and striatum. Scale bar = 1 mm. **g**. Quantification of % LFB+ area ((LFB area / ROI area) * 100) in the corpus callosum and striatum. Mean +/− SEM; n = 3 mice/genotype. *p < 0.05 (*), two-way ANOVA. **h**. Transmission electron microscopy (TEM) of myelin ultrastructure in the cortex, corpus callosum, and striatum from female (top) and male (bottom) *Nes*^Cre^hy*TAF1*^XDP^ brains. Scale bars = 1 µm. **i**. Representative TEM images highlighting pathological myelin phenotypes in *Nes*^Cre^hy*TAF1*^XDP^ brains. Yellow arrows indicate myelin splitting, ballooning, and abnormal ensheathment of multiple axons. Examples shown from both female (top) and male (bottom) brains. Scale bars = 1 µm.

Next, we assessed the changes to OL-lineage cells. *Nes*^Cre^hy*TAF1*^XDP^ had significantly lower proportion of SOX10+ cells in the corpus callosum compared to controls (Figure 5b,d). This could be because the corpus callosum is the hub or center of OL-lineage cells which actively make mature OLs, and this process may be disrupted in XDP. We also compared the percentage of PDGFRA+ oligodendrocyte precursor cells (OPCs) in motor cortex, corpus callosum, and striatum between *Nes*^Cre^ control and *Nes*^Cre^hy*TAF1*^XDP^ genotypes and found significant increase in PDGFRA+ cells proportion in the striatum of *Nes*^Cre^hy*TAF1*^XDP^ mice. No significant differences were detected in the motor cortex or corpus callosum (Figure 5c,e).

Using Luxol Fast Blue (a histological stain that identifies myelin), we visualized myelin-associated phospholipids and neural structures in both *Nes*^Cre^ control and *Nes*^Cre^hy*TAF1*^XDP^ mouse brains (Figure 5f). We observed a significant reduction in myelin lipids in the corpus callosum of *Nes*^Cre^hy*TAF1*^X-DP^ mice further supporting impaired myelination in this area (Figure 5g).

To examine the ultrastructural details of this disrupted myelination, we performed transmission electron microscopy (TEM). For comparison, we included female *Nes*^Cre^hy*TAF1*^XDP^ mice, as they are heterozygous carriers of mutant hy*TAF1* and can provide insight into whether the presence of a single functional *Taf1* allele is sufficient to maintain normal myelination. Strikingly, while female *Nes*^Cre^hy*TAF1*^XDP^ brains displayed relatively intact myelin structures in the cortex, corpus callosum, and striatum, male *Nes*^Cre^hy*TAF1*^XDP^ mice showed pronounced disruption of fine myelin architecture in all three regions (Figure 5h). Specifically, in the corpus callosum, we observed an almost complete loss of dense, compact myelin in *Nes*^Cre^hy*TAF1*^XDP^ male brains compared to females (Supplementary Figure S7c). Furthermore, in the female striatum, myelin was organized into fibrous tracts characteristic of the region’s striated structure. In males, however, myelin appeared as sparse, compact pockets rather than continuous fibrous tracts (Supplementary Figure S7d). Overall, compared to females, myelin in male *Nes*^Cre^hy*TAF1*^XDP^ brains was structurally disorganized, less compact, and exhibited thinner sheaths, indicating impaired myelin integrity.

We further characterized the ultrastructural myelin abnormalities observed in both female and male *Nes*^Cre^hy*TAF1*^XDP^ brains. In heterozygous *Nes*^Cre^hy*TAF1*^XDP^ females, we observed atypical features such as myelination of double axons (likely a result of axonal sprouting)^23^ and splitting of myelin lamellae, which may indicate axon-myelin uncoupling and ballooning or vacuolation, suggestive of impaired myelin integrity (Figure 5i). Since *TAF1* is located on the X chromosome, female mice are mosaics due to X chromosome inactivation, with individual cells expressing either the WT or XDP allele. It is possible that the majority of cells in our analysis had inactivated the XDP allele, while cells that inactivated the WT allele (and therefore expressed the mutant XDP allele) were more vulnerable and died prior to analysis. In male *Nes*^Cre^hy*TAF1*^XDP^ brains, these abnormalities were more pronounced, with significantly disrupted myelin ultrastructure indicative of axonal sprouting and outfoldings^24,25^. Notably, myelin outfoldings are a pathological feature in certain neuropathies and can be linked to disruptions in myelin synthesis, disruption in cytoskeletal dynamics, and the improper organization of myelin membranes^26,27^. Additionally, male brains showed many more split myelin sheaths and ballooned structures, consistent with severe myelin pathology (Figure 5i).

### Human XDP patients have myelin loss in the medial prefrontal cortex and aberrant myelination in the striatum

A previous study using quantitative MRI in a small cohort of XDP patients suggested potential involvement of white matter underlying XDP pathology^28^. However, myelin alterations and the associated OL cellular changes in XDP brains have not been previously defined. To investigate this and validate our findings from the XDP mouse model in humans, we obtained postmortem brain tissue from XDP patients and age-matched healthy controls to assess myelin integrity. We first focused on three brain regions: the ventral division of the medial prefrontal cortex (Brodmann area 24; “MFC”), the rostral portion of the corpus callosum (“CC”), and the cingulum bundle within the cingulate cortex (“Cg Ctx”) (Figure 6a, Supplementary Figure S8a). Luxol Fast Blue staining revealed a selective reduction of myelin in the MFC of XDP patient brains (Figure 6b).

**Figure 5.**
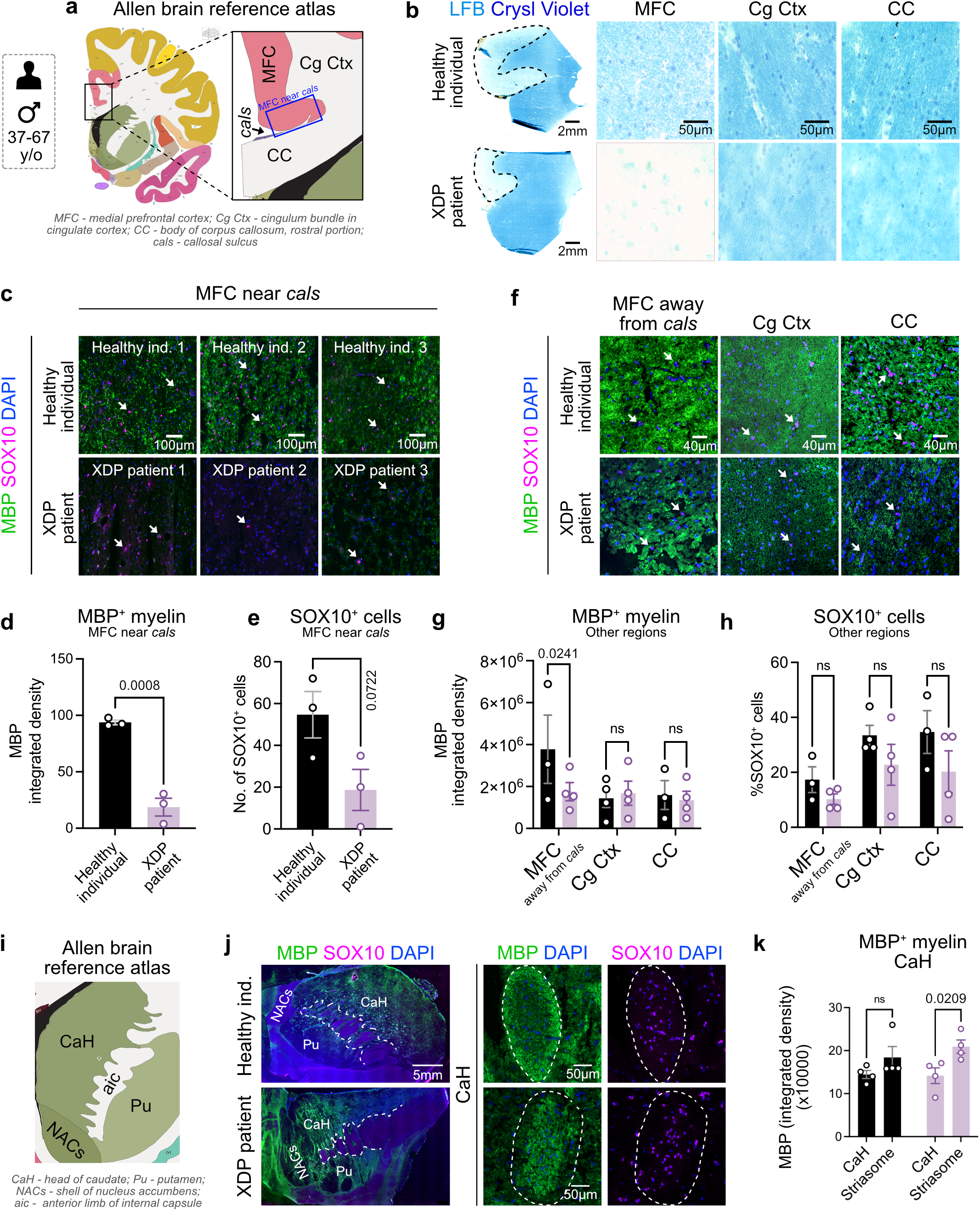
Myelin pathology in postmortem brain tissue from XDP patients. a. Schematic from the Allen Brain Atlas highlighting brain regions sampled from 40-63 year old male XDP patients and age-matched healthy controls. Regions include the medial frontal cortex (MFC), cingulum bundle in the cingulate cortex (Cg Ctx), rostral corpus callosum (CC), and callosal sulcus (cals). The blue box indicates the MFC area near the callosal sulcus. **b**. Luxol Fast Blue (LFB) staining of myelin lipids in large brain sections and subregions (MFC, Cg Ctx, CC) from XDP and control brains. XDP samples show reduced LFB staining in the MFC. Scale bar = 50 µm. **c**. Immunostaining of the MFC near cals for myelin basic protein (MBP, green) and oligo-dendrocyte lineage marker SOX10 (magenta) reveals reduced MBP signal in XDP brains. White arrows indicate SOX10+ cells. Scale bar = 100 µm. **d**. Quantification of MBP+ myelin (integrated density = fluorescence intensity × area) in the MFC near cals. Healthy ind. = healthy individual. Data are shown as mean +/− SEM; n = 3 individuals per group. Unpaired t-test; p = 0.0008. **e**. Quantification of SOX10+ cells (% SOX10+ = SOX10+/DAPI+ × 100) in the same region. Mean +/− SEM; n = 3 per group. Unpaired t-test; p = 0.0722. **f**. Immunostaining of MFC (away from cals), Cg Ctx, and CC for MBP and SOX10 in XDP brains. White arrows indicate SOX10+ cells. Scale bar = 100 µm. **g**. Quantification of MBP+ myelin in MFC (away from cals), Cg Ctx, and CC. Mean +/− SEM; n = 3 individuals per group. ns: non-significant, two-way ANOVA. **h**. Quantification of SOX10+ cells in MFC (away from cals), Cg Ctx, and CC. Mean +/− SEM; n = 3 individuals per group. ns: non-significant, two-way ANOVA. **i**. Schematic from the Allen Atlas showing dissected striatal regions including the head of the caudate (CaH), putamen (Pu), nucleus accumbens shell (NACs), and anterior limb of internal capsule (aic). **j**. Immunostaining of the CaH for MBP and SOX10 reveals reduced myelin signal in XDP brains. White arrows indicate SOX10+ cells. Healthy ind. = healthy individual. Scale bars: whole section = 1 mm; insets = 50 µm. **k**. Quantification of MBP+ myelin in the CaH. Mean +/− SEM; n = 3 individuals per group. P value on plot, ns: non-significant, two-way ANOVA.

We next conducted a more detailed analysis of the MFC and observed striking differences in myelination across its subregions. In XDP patient brains, the region located immediately adjacent to the callosal sulcus, which is a boundary along the corpus callosum (Figure 6a, blue box), exhibited a near-complete loss of MBP compared to the same region in the brains of healthy individuals (Figure 6c-d, Supplementary Figure S8b). Quantification revealed a significant reduction in MBP+ myelin in all three independent XDP cases relative to three independent healthy controls within MFC (Figure 6c-d, Supplementary Figure S8c). There was no significant difference in the number of SOX10+ cells in this area (Figure 6c-e, Supplementary Figure S8b).

We also examined MBP+ myelin in additional regions of the MFC (distant from the callosal sulcus), as well as in other brain areas, including the Cg Ctx and the CC (Figure 6f). In XDP patient brains, MBP+ myelin was significantly reduced in the MFC region distal to the callosal sulcus compared to healthy individuals, whereas no significant differences were observed in the Cg Ctx or CC (Figure 6f-g, Supplementary Figure S8d). The %SOX10+ cells also did not significantly change in other areas of the MFC, Cg Ctx, or the CC (Figure 6f-h, Supplementary Figure S8e).

We next analyzed striatal tissue from XDP patients and healthy individuals (Figure 6i). Consistent with our observations in *Nes*^Cre^hy*TAF1*^XDP^ male mice where MBP accumulated within the core of the striosomes rather than displaying the typical halo-like pattern along their borders (Figure 5a, Supplementary Figure S7a), we observed a similar distribution of MBP+ myelin in the striatum of XDP patient brains (Figure 6j,k). The %SOX10+ cells was not significantly different in head of caudate (“CaH”) region of XDP patient versus healthy individual brains (Supplementary Figure S8e-f). These data suggest selective myelin disruption is most likely caused by oligodendrocyte dysfunction within the MFC and striatum to be a key feature of the human XDP brain.

## DISCUSSION

Along with our companion paper^18^, we introduce a novel humanized mouse model of XDP, a neurodegenerative movement disorder caused by the insertion of a hominoid-specific SINE-VNTR-*Alu* (SVA) retrotransposon within intron 32 of the *TAF1* gene. This model features a conditional hybrid mouse-human *Taf1*/*TAF1* allele that includes the XDP-specific pathogenic intronic SVA_F element. By enabling Cre-dependent expression of this hybrid gene, we can direct expression of the disease allele to specific cell types. In this study, we induced the conversion of hy*TAF1* in Nestin+ neural progenitor cells, which gave rise to male *Nes*^Cre^hy*TAF1*^XDP^ mice exhibiting phenotypes that closely mirror clinical manifestations observed in human XDP patients.

Male *Nes*^Cre^hy*TAF1*^XDP^ mice show markedly reduced body size and weight compared to their *Nes*^Cre^ control littermates and also have a shortened lifespan. Despite their shorter lifespan, we successfully performed longitudinal tracking of motor behaviors from P14 through P35. During this window, *Nes*^Cre^hy*TAF1*^XDP^ males demonstrated progressive motor function deficits relative to both male *Nes*^Cre^ controls and heterozygous female carriers, consistent with the sex-specific penetrance of XDP. Specifically, these males exhibited reduced motor coordination and balance, as assessed by rotarod and pole test performance. Additionally, they showed impaired fore and hind limb strength in grip strength tests and characteristic gait abnormalities, including decreased swing speed, reduced duty cycle, and altered paw position. These motor phenotypes recapitulate clinical features of XDP and offer a valuable behavioral readout for preclinical therapeutic testing.

A key neuropathological finding in this model is the down-regulation of OL-specific genes and associated disruptions in myelin structure. OLs, the myelinating glial cells of the CNS^29^, are increasingly recognized as contributors to various neurodegenerative conditions, including multiple sclerosis^30,31^, leukodystrophies^32^, Alzheimer’s disease^33^, and Parkinson’s disease^34^. In our model, bulk RNA sequencing revealed robust downregulation of genes critical for the function of mature, myelinating OLs across both the cortex and striatum. In contrast, transcript levels of oligodendrocyte precursor cells (OPC) markers^35^ such as *Cspg4* (encoding NG2) and *Pdgfra*^36,37^ remained unchanged, indicating that the OPC population may not be depleted at the transcript level, at least during the stage of disease examined. Immunohistochemical analysis revealed a significant reduction in the percentage of SOX10+ cells in the corpus callosum, a white matter-rich region densely populated by both OPCs and mature OLs. As SOX10 is expressed across the OL lineage, including OPCs and newly differentiating OLs, this reduction may reflect a broader impairment in OL homeostasis, either due to cell loss or altered differentiation capacity.

We observed a significant increase in the proportion of PDGFRA+ cells in the striatum of *Nes*^Cre^hy*TAF1*^XDP^ male mice. This could represent an expansion or accumulation of OPCs in response to injury or myelin disruption. Similar increases in OPC proliferation have been reported in the striatum of Huntington’s disease models^38^, where they are thought to represent a compensatory mechanism aimed at replenishing the mature OL pool and restoring myelin integrity. However, whether the expanded OPC pool in XDP is functional and capable of differentiating into mature, myelinating OLs remains unclear. Taken together, these findings suggest that OL lineage cells are dysregulated in XDP in a regionally specific manner, which could contribute to potential failures in maintaining mature OLs or in generating them from precursors. Whether disease progression is primarily driven by the degeneration of existing mature OLs, impaired differentiation of OPCs into OLs, or a combination of both remains to be explored.

Importantly, these myelin abnormalities are also found in human XDP patients. Postmortem analysis of human XDP brain tissue revealed similar myelin pathology. In the medial prefrontal cortex, regions adjacent to the callosal sulcus showed near-complete loss of myelin compared to brains of healthy individuals. In the striatum, MBP+ myelin was abnormally distributed within the striasomes, accumulating centrally rather than along the borders. A strikingly similar distribution of myelination was also observed in our XDP mice. These findings support a conserved pathological feature of aberrant myelination in both human patients and XDP mice. Currently, there are no disease-modifying therapies for XDP, and treatment remains limited to symptomatic management. While deep brain stimulation of the globus pallidus interna can alleviate dystonia in some patients, it does not slow down disease progression. Neuroimaging studies, including diffusion tensor imaging, have indicated compromised white matter integrity and associated axonal tract disruptions in the striatum^12,28,39^. Extremely limited data on tissue pathology particularly during pre-symptomatic stages has made it challenging to track the cellular and molecular changes underlying white matter alterations across different brain regions in patient tissue.

Our study highlights OL dysfunction and myelin disruption as central pathological features of XDP. This model opens new avenues for exploring the cell-type-specific mechanisms underlying disease onset and progression. Future studies will define whether OL dysfunction is a cause or consequence of neurodegeneration in XDP. By dissecting the temporal and functional contributions of OLs, we can begin to fill critical gaps in our understanding of XDP pathogenesis. Importantly, these insights are likely to have broader relevance beyond XDP. Both Parkinson’s disease and Huntington’s disease are characterized by striatal pathology and motor dysfunction, and emerging evidence suggests that OL dysfunction and white matter changes may contribute to disease mechanisms in these disorders as well. Our findings therefore not only deepen our understanding of XDP but may also provide a valuable framework for investigating shared glial contributions to neurodegeneration across multiple disorders.

## METHODS

### Animals

All animal experiments were approved by NYU Grossman School of Medicine’s institutional Animal Care and Use Committee (IACUC). Nestin^Cre^ (*Nes*^Cre^) mice (B6.Cg-Tg(Nes-Cre)1Kln/J) were purchased from Jackson Laboratories (Strain #003771). Mice were housed on a 12 h light/ dark cycle and were given food and water ad libitum. All animal procedures were in accordance with the guidelines provided by the National Institute of Health as well as NYU School of Medicine’s Administrative Panel on Laboratory Animal Care. All animals were housed at 22-25 °C and 50-60% humidity.

### Human tissue

Human XDP post-mortem brain tissue was obtained from Collaborative Center for X-linked Dystonia Parkinsonism (CCXDP)^40^. The clinical characterization of patient donors including age of onset (50.8 ± 5.14 years non-symptomatic control; 52.25 ± 4.05 years XDP), initial symptoms and symptoms at the time of death, brain weight (616 ± 18.3 g non-symptomatic control; 582.5 ± 2.5 g XDP), cause of death, hexameric repeat length, etc., were assessed by clinical experts. Postmortem interval was also noted (21.6 ± .9 h non-symptomatic control; 30.2 ± 11.3 h XDP) but not controlled for.

### Phenotype assessment

#### Body weight

These data were analyzed by fitting a mixed-effects model to handle the missing values. Each row in the dataset represents a different time point, so matched values are stacked into a sub-column. The analysis did not include an interaction term (fit a main effects only model i.e., column effect (genotype) and row effect (time) only. Sphericity was assumed (i.e., no correction was applied). ****p<0.0001 and *p=0.0179.

#### Behavior

##### Husbandry

Three-week old mice were transferred to the satellite housing facility of the Rodent Behavior Laboratory at NYU Langone Health immediately after weaning. Since the mice were slightly underdeveloped, hydrogel was provided to all cages from week 3 to 5. On the first week after weaning, regular chow was slightly wetted and placed on the floor of the cage to ensure access to food. During the behavioral testing period, the animals were weighed daily and closely monitored for any signs of distress or health problems. Each behavioral test was performed once a week between week 3-5, on different days to limit fatigue.

##### Rotarod

The rotarod test is widely used to measure locomotor function, balance, coordination and learning. Rotarod performance was assessed by how long animals were able to run on a rotating cylindrical rod of increasing speed (Model 755, IITC Life Science). The standard protocol of the NYU Rodent Behavior Laboratory was as previously published^41,42^. Individual mice were first placed on the horizontal, static rod (3 cm diameter, 30 cm above a padded floor base) for a minimum duration of 60 s to habituate the animal to handling and the apparatus. On all subsequent trials, each mouse was placed onto the rod for at least 10 s before initiating the rotation program with revolutions per min (rpm) set to ramp in a linearly accelerating manner from 4 to 40 rpm over a 5 min trial. Each mouse was tested for 6 trials and allowed to rest in their home cage for at least 25 min between trials. The latency to fall from the rod was recorded and statistically analyzed, with body weight included as a covariate. Data from trials when an animal could not stay on the static rod for at least 10 s prior to rotation (given 3 attempts) were marked as failed and excluded.

##### Vertical pole test

The vertical pole test assesses the motor ability of mice to grasp and maneuver their limbs and body to descend a pole^43-45^. The pole comprised a plain steel rod (50 cm height and 1.25 cm diameter) wrapped in paper tape. The pole was vertically oriented with its base cushioned with polypropylene fabric and the top capped with a 15 cm diameter plastic disk to prevent climbing onto the tip. On each trial, an individual mouse was positioned to grasp the pole at the top facing upwards and given a maximum of 90 s to turn and descend the pole to its base. On week 3, mice were assessed on a session of 8 trials whereas 6-trial sessions were performed on later timepoints. A minimum of 20 min was given between trials for the same mouse. The latency to turn to orient downward and the total time to descend the pole (from initial pole placement to reaching the base with all four paws) were recorded and confirmed by video analysis from a side-view camera. Mean scores on each measure across the last six trials were compiled for statistical analysis, with body weight included as a covariate. Data from trials when the mouse couldn’t grasp the pole, or fell from it, were marked as failed and excluded.

##### Grip strength

Limb and digit grip strength was assessed as described previously^41,42^. A sensitive force sensor apparatus (BIO-GS3, Bioseb) was used to quantify the horizontal force generated by each mouse while grasping a 6x10 cm stainless steel grid platform. Two different grip strength indices were collected: forelimb only and combined fore- and hind-limbs (all-limb). Individual mice were subjected to 6 trials on each index with a minimum inter-trial interval (ITI) of 30 s and at least 30 min between forelimb and all-limb assessment. On forelimb grip trials, each mouse was positioned so that its forepaws grasped the central top-half portion of the grid. The mouse was then pulled away from the grid by the base of its tail until their grip was released, ensuring that the torso remained in a horizontal position. All-limb grip trials were performed similarly, except that all four paws of the mouse were placed centrally on the grid. The placement of the animal on the platform, pulling force, angle, and speed were kept as consistent as possible to minimize variability. The maximum force achieved (in grams) on each trial was recorded from the digital display of the grip strength meter. For both forelimb and all-limb measures, the truncated mean of the 6 trials (highest and lowest scores removed) were statistically analyzed, with body weight included as a covariate. Data from trials when the mouse were unable to grasp the grid were marked as failed and excluded.

##### Gait analysis

Gait was assessed through quantitative analysis of footfalls and parameters of quadrupedal locomotion in freely moving animals using the Catwalk XT (software version 10.7.704, Noldus Information Technology). For each trial, individual mice were placed on one end of the enclosed runway (200 cm length, 10 cm width, 20 cm height) and allowed to traverse unrestrained at their own pace. Mice were allowed to explore the runway until they generated compliant runs; in which they were required to traverse the middle 60 cm of the runway at a minimum velocity of 5 cm/s and 20 footfalls within 10 s, without long stalls or alternate behaviors. In each session, a mouse was run until it achieved a minimum of 3 compliant runs. A high-speed camera located underneath the transversely illuminated glass runway captured detailed data for each footfall. All footfalls were validated manually after automated classification. Detailed gait parameters for each mouse were then automatically computed and extracted based on the digital paw prints acquired during the runs. Body weight and average speed were included as covariates in the statistical analysis whenever they exerted a significant impact on the outcome measure of interest.

##### Statistical testing

Statistical analysis of the motor behavior data was analyzed using linear mixed effects models with age/timepoint as within-subjects and genotype and sex as between-subjects fixed factors. The repeated measures within-subjects covariance matrix was modeled using an unstructured covariance structure. Body weight was included as a time-varying covariate for when analyzing measures in the vertical pole, grip strength and rotarod. For gait analysis, body weight and/ or speed were included as time-varying covariates whenever contributing significantly to the mixed effects model. All statistical analyses were performed using IBM SPSS Statistics (version 28.0.0.1)

### Bulk RNA sequencing

#### Tissue and RNA processing

Brain tissues were homogenized using a disposable Pellet Pestle (Fisher Scientific, 12-141-364) in 350 μl of buffer RLT (QIAGEN, 74136). Total RNA was extracted from mouse brain tissue using the Qiagen RNeasy Plus Mini Kit according to the manufacturer’s protocol, including on-column DNase digestion to eliminate genomic DNA contamination.

RNA integrity was assessed using the Agilent 2100 Bioanalyzer, and only samples with RNA integrity numbers (RIN) ≥ 8.0 were used for sequencing. Bulk RNA-seq libraries were constructed using Illumina’s TruSeq Stranded mRNA Library Prep Kit, starting with 200 ng of total RNA per sample and amplifying the libraries with 12 cycles of PCR. Final libraries were visualized using High Sensitivity DNA ScreenTape on the Agilent TapeStation 2200 instrument. Quant-It was used for final concentration determination and libraries were pooled equimolar. Sequencing was carried out on an Illumina NovaSeq+ platform with paired-end 50 bp reads, generating approximately 30 million reads per sample.

#### RNA-Seq data analysis

Raw reads were processed using the Seq-N-Side RNA-STAR pipeline^46^. Adapter sequences and low-quality bases (Phred score < 20) were trimmed using Trimmomatic (v0.36)^47^. Read quality was assessed using FastQC before and after trimming. High-quality reads were aligned to the mouse reference genome (GRCm38/mm10) using STAR (v2.7.3a)^48^, with splice junctions guided by the GENCODE M25 GTF annotation. STAR was run with standard parameters for spliced alignment, retaining only uniquely mapped reads. The mean read insert sizes, and their standard deviations were calculated using Picard tools (v.2.18.20) (http://broadinstitute.github.io/picard). The genes-samples counts matrix was generated using featureCounts (v1.6.3)^49^, normalized based on their library size factors using DEseq2(v1.30.1)^50^, and differential expression analysis was performed. The Read Per Million (RPM) normalized BigWig files were generated using deepTools(v.3.1.0)^51^. To compare the level of similarity among the samples and their replicates, we used two methods: principal-component analysis and Euclidean distance-based sample clustering. All the downstream statistical analyses and generating plots were performed in R environment (4.0.3).

### Pathology

#### H&E staining

Mounted, fresh frozen mouse brain tissue (16 µm coronal sections) was taken from storage at −80 ºC and thawed to 37 ºC for 5 min. Then, the slides were incubated at −20 ºC in chilled methanol for 30 min. After incubation, the slides were transferred to ethanol for 1 min at room temperature (RT). The slides were left to air dry for no longer than 10 min, and then were treated with Hematoxylin (Agilent Technologies, S330930-2) for 7 min at RT. After removing the hematoxylin solution, the slides were rinsed 5 times with ultrapure water, then rinsed 15 times in a separate water beaker, and another 15 times in a 3rd water beaker. The slides were then treated with Bluing Buffer solution (Agilent Technologies, CS70230-2) for 2 min at RT. After Bluing Buffer was removed, the slides were rinsed 5 times with ultrapure water. Then slides were treated with Eosin Mix (10% Eosin Y Solution (Sigma-Aldrich, HT110216) in Tris-Acetic Acid Buffer [0.45 M, pH 6.0]) for 1 min, and rinsed 15 times in ultrapure water. After, they were placed on a slide warmer at 37 ºC for 5 min and mounted with Fluoromount (SouthernBiotech, 0100-01) before imaging on KEYENCE microscope.

#### Luxol Fast Blue staining

Mounted, fresh frozen brain tissue (16 µm coronal sections) was taken from storage at −80 ºC and rehydrated in 95% ethyl alcohol for 2 min at room temperature (RT). Slides were then placed in 0.1% Luxol fast blue solution (0.1 gm Luxol fast blue, MBS (VWR, TS21217-0250), 100 mL 95% ethyl alcohol, 0.5 mL glacial acetic acid) and left in a 56 ºC incubator overnight (about 16 h). After removing from Luxol fast blue solution, excess stain was rinsed off with 95% ethyl alcohol and slides were rinsed in distilled H2O. Slides were then differentiated in 0.05% lithium carbonate solution (0.05 gm lithium carbonate (Sigma-Aldrich, 255823), 100 mL distilled H_2_O) for 30 s and then in 70% ethyl alcohol for 30 s, then rinsed in distilled H_2_O. This differentiation process was repeated 2 times total, until grey matter appeared clear and white matter was sharply defined under a microscope. After differentiating, slides were counterstained in 0.1% Cresyl violet solution (Abcam, ab246816) for 30-40 s, then rinsed in distilled H_2_O. Slides were differentiated in 95% ethyl alcohol for 5 min, and then placed in 100% alcohol 2 times, for 5 min each. Then (working in a fume hood) slides were placed in xylene 2 times, for 5 min each. Slides were then mounted with EcoMount (Biocare Medical, EM897L) and imaged on KEYENCE microscope.

#### Immunofluorescence staining

Mounted, fresh frozen brain tissue (16 µm coronal sections) were taken from storage at −80 ºC and post-fixed in 4% PFA at room temperature (RT) for 15-20 mins. Slides were washed 3 times in PBS, 5 min each. If necessary, antigen retrieval was then performed by submerging slides into boiling citrate buffer, pH = 6.0 (Sigma-Aldrich, C9999) for 10 min. Slides were blocked for 1 h at RT with a blocking buffer solution consisting of 0.4% Triton X-100 (Sigma-Aldrich, T8787) and 10% normal goat serum or normal donkey serum depending on 2º Ab host species. After blocking, slides were left to incubate in 1º Ab solution in a 4 ºC humidified chamber overnight. The following day the 1º Ab solution was drained off and slides were washed 3 times in PBS, 5 min each. Slides were then incubated with the appropriate secondary antibodies diluted 1:300 in blocking buffer solution for 90 min at RT. After 2º Ab incubation, slides were again washed 3 times in PBS for 5 min each. Slides were then covered with 0.001% DAPI solution (in PBS) for 3 min, and then with 5% TrueBlack (Biotium, 23007) diluted in 70% EtOH for 30 s. Slides were rinsed 4 times in PBS to wash off any excess dye, then mounted using Fluoromount (SouthernBiotech, 0100-01) and imaged on either an LSM 800 confocal or KEYENCE microscope.

#### Electron microscopy

Anesthetized mice were perfused transcardially with saline followed by the fixative containing 4% paraformaldehyde, 2.5% glutaraldehyde and 0.1 M sucrose in 0.1 M sodium cacodylate buffer (CB, pH 7.4). After perfusion, the mouse was covered with wet paper towel and fixed in situ for 2 h. Mouse brain was dissected, sagittal cut from the middle, and 1 mm brain slice from one side of the brain was obtained for analysis of corpus callosum. The other half brain was dissected coronal for the region of stratum and motor cortex. All dissected samples were continue fixed in the same fixative at 4 °C overnight. Fixed tissues were washed with CB 3 x 15 min, post-fixed with 2% osmium tetroxide in CB for 2 h, dehydrated in the serial of ethanol solutions (30, 50, 70, 85, 95, 100%, 100%; 15 min each), then infiltrated with propylene oxide and embedded in EMbed 812 (Electron Microscopy Sciences, Hatfield, PA). 1 µm semi-thin section was cut and stained with toluidine blue to evaluate the region of the interests. Ultrathin sections (80 nm) were cut, mounted on formvar coated slot copper grids and stained with uranyl acetate and lead citrate. Stained grids were imaged under Zeiss Gemini300 FESEM using the STEM detector for the large field of view, and JEOL 1400 Flash transmission electron microscope (Japan) with Gatan Rio16 camera (Gatan, Inc., Pleasanton, CA) for the high-resolution imaging.

#### Stastics and reproducibility

Except for specific statistical analysis reported elsewhere, statistical analyses were performed using GraphPad Prism (v9.0) or R for all datasets except behavioral experiments, which were analyzed using IBM SPSS Statistics (v27). Unpaired two-tailed Student’s t-tests were used for comparisons between two groups. For experiments involving multiple groups or repeated measures, two-way ANOVA with Tukey’s or Šídák’s post hoc test was used as appropriate. Behavioral data were analyzed in SPSS using mixed-model ANOVA to account for within-subject factors (time point) and between-subject variables (genotype). Statistical significance was defined as p < 0.05. Data are shown as mean ± standard error of the mean unless otherwise indicated. The specific significant p values and sample sizes (number of mice per genotype) are indicated in each figure and/or in the figure legend.

## ACKNOWLEDGEMENTS

This work was funded by the Collaborative Center for X-linked Dystonia Parkinsonism (S.A.L., J.D.B.), MD Anderson Neurodegeneration Consortium (S.A.L.), Anonymous Donors, the Carol and Gene Ludwig Family Foundation (S.A.L.). We thank the Leon Levy Foundation for postdoctoral fellowship in neuroscience award (P.P.). Behavioral experiments were performed in the NYULMC Rodent Behavior Laboratory (RRID: SCR_017942) supported by the NIH BRAIN Initiative U19NS1076 (A.C.M.). We thank NYU Langone Health’s Genome Technology Center (RRID: SCR_017929). This shared resource is partially supported by the Cancer Center Support Grant P30CA016087 at the Laura and Isaac Perlmutter Cancer Center. We also thank Aristotelis Tsirigos and Ziyan Lin at NYU Langone Health’s Applied Bioinformatics Laboratories (RRID:SCR_019178) for assistance with bulk RNA sequencing data analysis. The Applied Bioinformatics Laboratories is partially supported by NYU Langone’s Laura and Isaac Perlmutter Cancer Center support grant P30CA016087. We thank NYU Langone Health DART Microscopy Lab personnel Alice Liang, Joseph Sall and Jason Liang for their consultation and assistance with EM work. This core is partially funded by NYU Cancer Center Support Grant NCI P30CA016087 and Gemini300 FESEM was supported by NIH S10 OD019974. We thank Drs. ngelina Evangelou and James Salzer for providing the SOX10, PGDFRA, and MBP antibodies. We thank Prof Klaus-Armin Nave and Dr Wiebke Möbius for their thoughtful discussion on the electron micrographs and myelin pathology. We thank the members of the Liddelow lab for their comments and discussions on the project. Select illustrations were made using BioRender.com.

## AUTHOR CONTRIBUTIONS

P.P. and S.A.L. conceived the study. P.P. designed and performed the experiments. W.Z., Y.Z., R.B., and J.D.B. generated the hyTAF1PC-XDP pre-converted mice. K.L. and A.O.K. assisted with mouse maintenance, breeding, and phenotyping assays. K.F.L. and A.C.M. contributed to behavioral experiments. K.L. also performed immunohistochemistry and imaging. H.A. assisted with mouse genotyping, W.Z. with RNA extraction, and M.R.O. with bulk RNA sequencing data analysis. C.F.C., G.P.A.L., M.S., E.L.M., M.A.C.A., C.C.E.D., J.H., E.N., E.B.P., and D.C.B. were involved in the acquisition and sectioning of human postmortem tissue. S.A.L., J.D.B., and R.B. supervised the study. S.A.L. and J.D.B. acquired funding. P.P. drafted the manuscript, and all authors reviewed and approved the final version.

---

**Supplemental Figure S1.**
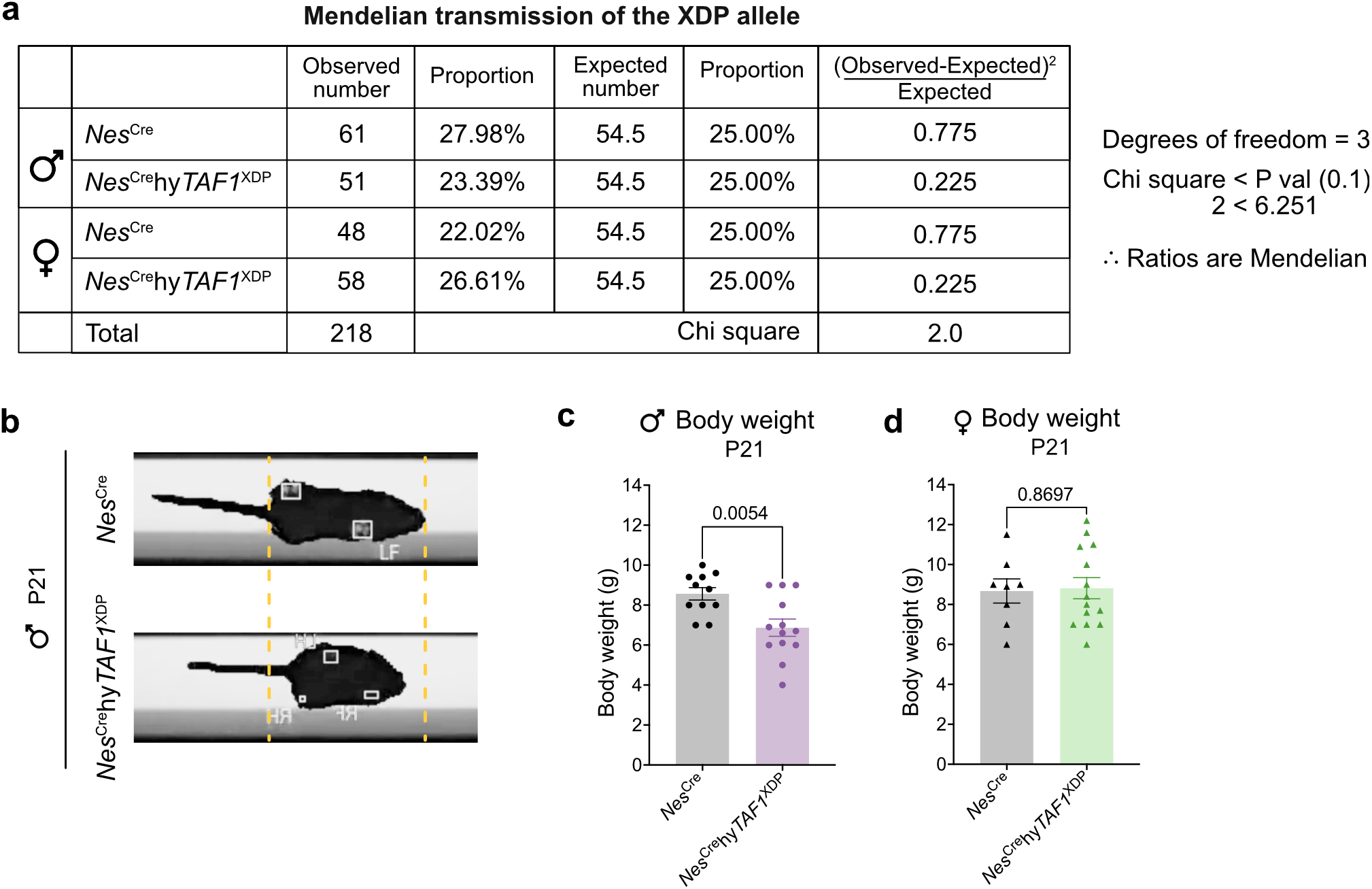
Mendelian inheritance and phenotype of *Nes*^Cre^ and *Nes*^Cre^hy*TAF1*^XDP^ mice. **a**. Mendelian ratio analysis of genotypes from a cross between *Nes*^Cre^ and hy*TAF1*^PC-XDP^ alleles. Observed numbers for four progeny genotypes were compared to expected Mendelian ratios (25% each) using a chi-square test. The observed proportions did not significantly deviate from expected values (χ^2^ = 2.0, df = 3, p > 0.1), indicating normal transmission of the humanized *Taf1* allele and no germline disruption from the insertion. **b**. Representative image comparing body size of male *Nes*^Cre^ and *Nes*^Cre^hy*TAF1*^XDP^ mice at P21. Yellow dotted line outlines the size of a *Nes*^Cre^ control male for visual reference. **c**. Quantification of body weight in male *Nes*^Cre^ and *Nes*^Cre^hy*TAF1*^XDP^ mice at P21. *Nes*^Cre^hy*TAF1*^XDP^ males exhibit significantly reduced body weight compared to controls. Unpaired t-test; p = 0.0054. Data shown as mean +/− SEM; n = 11 *Nes*^Cre^ and n = 13 *Nes*^Cre^hy*TAF1*^XDP^ mice. **d**. Quantification of body weight in female *Nes*^Cre^ and *Nes*^Cre^hy-*TAF1*^XDP^ mice at P21. No significant difference observed. Unpaired t-test; p = 0.8697 (ns). Mean +/− SEM; n = 8 *Nes*^Cre^ and n = 14 *Nes*^Cre^hy*TAF1*^XDP^ mice.

**Supplemental Figure S2.**
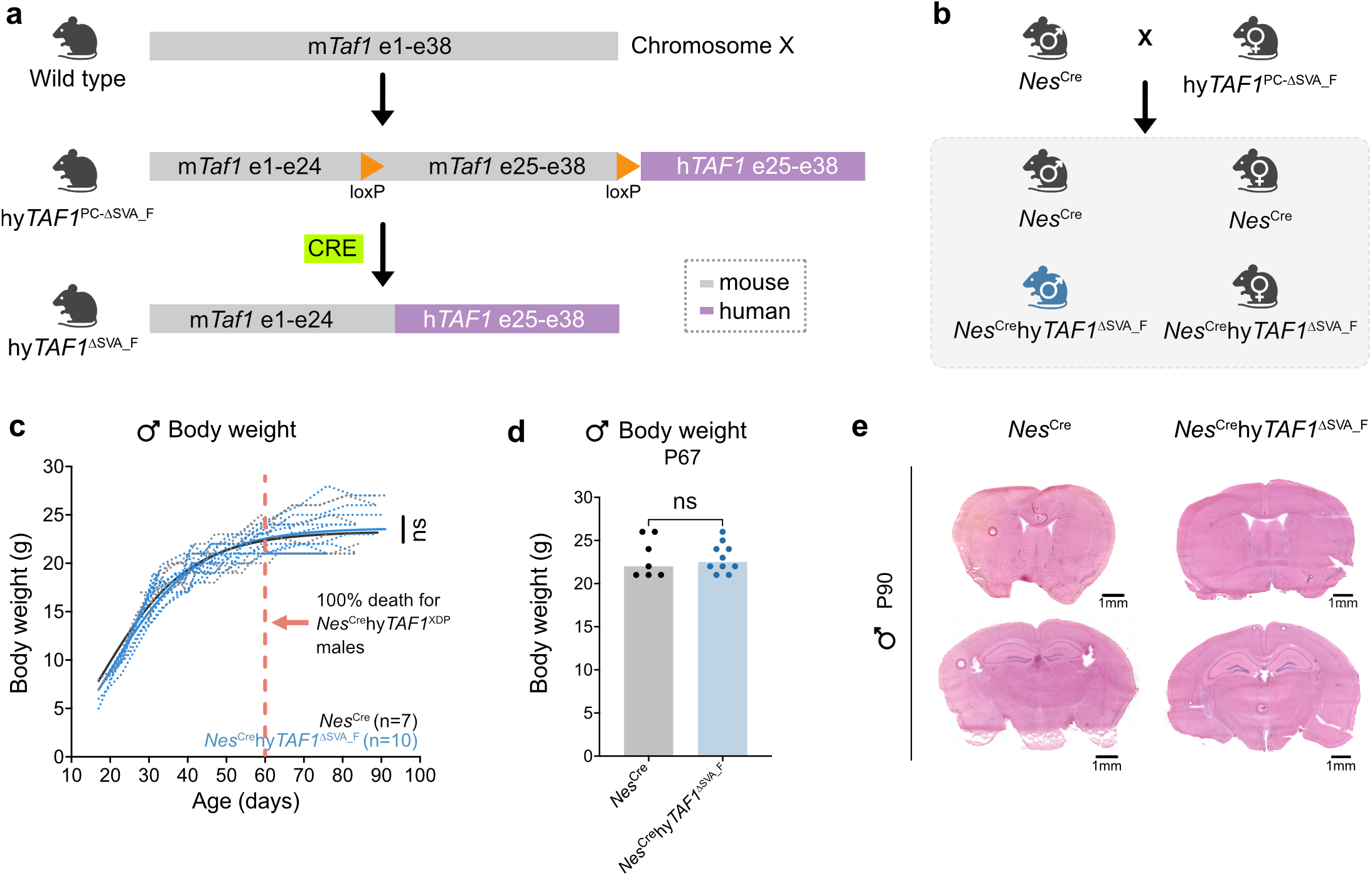
Generation and phenotypic characterization of *Nes*^Cre^hy*TAF1*^ΔSVA_F^ mice. **a**. Schematic illustrating the genetic engineering strategy for generating the hy*TAF1*^ΔSVA_F^ allele. (i) Wild-type mouse *Taf1* gene (m*Taf1*) spans exons 1-38 on the X chromosome (gray). (ii) An XDP patient-derived human *TAF1* segment (h*TAF1*; purple), encompassing exons 25-38 and lacking the disease-specific SVA_F retrotransposon, is inserted downstream of the mouse *Taf1* locus. (iii) *Nes*^Cre^-mediated excision of the m*Taf1* region results in a hybrid *TAF1* gene composed of exons 1-24 of mouse *Taf1* and exons 25-38 of human *TAF1*, thus generating an SVA_F-deleted humanized allele (hy*TAF1*^ΔSVA_F^). **b**. Breeding scheme: Crossing *Nes*^Cre^ homozygous males with heterozygous *Taf1*^PC-ΔSVA_F^ females produces four genotypes: *Nes*^Cre^ control males, *Nes*^Cre^hy*TAF1*^ΔSVA_F^ males (highlighted in blue), *Nes*^Cre^ control females, and *Nes*^Cre^hy*TAF1*^ΔSVA_F^ carrier females. **c**. Body weight trajectory of male *Nes*^Cre^ (n = 7) and *Nes*^Cre^hy*TAF1*^ΔSVA_F^ (n = 10) mice over time. Individual mice are shown with dotted lines; group means with bold lines. Statistical comparison via two-way ANOVA. **d**. Body weight at P67 of *Nes*^Cre^ and *Nes*^Cre^hy*TAF1*^ΔSVA_F^ males. Unpaired t-test. Data shown as mean +/− SEM. **e**. Representative H&E-stained coronal brain sections from P90 male *Nes*^Cre^ and *Nes*^Cre^hy*TAF1*^ΔSVA_F^ mice showing no gross anatomical abnormalities. Scale bar = 1 mm.

**Supplemental Figure S3.**
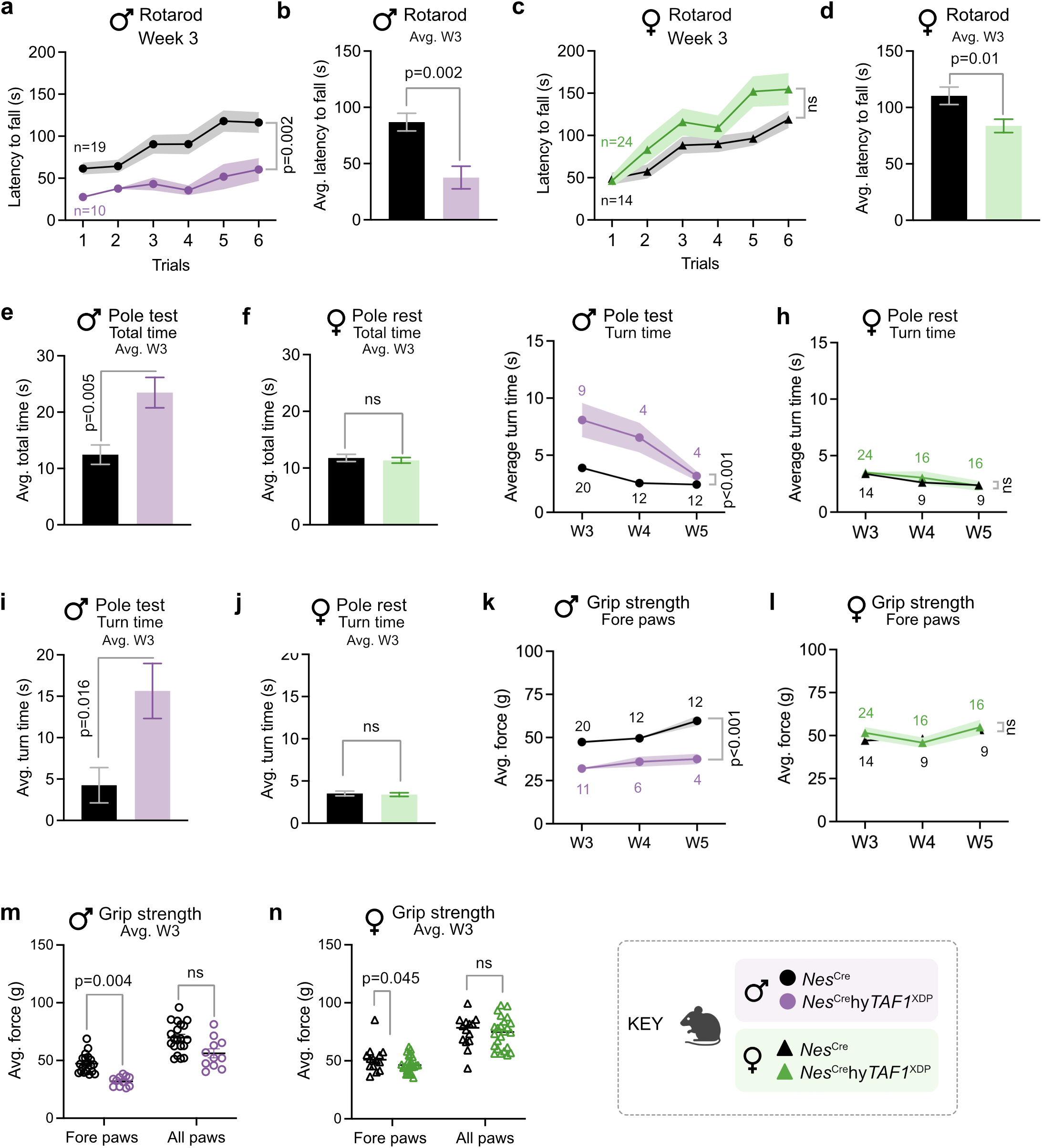
Rotarod, pole test, and grip strength analysis of *Nes*^Cre^hy*TAF1*^XDP^ male and female mice. **a**. Latency to fall (s) on the rotarod across trials in *Nes*^Cre^ and *Nes*^Cre^hy*TAF1*^XDP^ male mice. Data are shown as mean +/− SEM. p=0.002, unpaired t-test. Number of mice are indicated on the plot. **b**. Average latency to fall (s) at week 3 between *Nes*^Cre^ and *Nes*^Cre^hy*TAF1*^XDP^ male mice. p=0.002, unpaired t-test. **c**. Latency to fall (s) on the rotarod across trials in *Nes*^Cre^ and *Nes*^Cre^hy*TAF1*^XDP^ female mice. Data are shown as mean +/− SEM. ns: non-significant, unpaired t-test. Number of mice are indicated on the plot. **d**. Average latency to fall (s) at week 3 between *Nes*^Cre^ and *Nes*^Cre^hy*TAF1*^XDP^ female mice. p=0.01, unpaired t-test. **e**. Average total descent time (s) for male *Nes*^Cre^ and *Nes*^Cre^hy*TAF1*^XDP^ mice in the pole test at week 3. p=0.005, unpaired t-test. **f**. Average total descent time (s) for male *Nes*^Cre^ and *Nes*^Cre^hy*TAF1*^XDP^ mice in the pole test at week 3. ns, non-significant, unpaired t-test. **g**. Average turn time (s) taken by *Nes*^Cre^ and *Nes*^Cre^hy*TAF1*^XDP^ male mice to orient downward on the pole. p < 0.001, linear mixed-effects model with age and timepoint as within-subject factors and genotype as a between-subject fixed effect. **h**. Average turn time (s) taken by *Nes*^Cre^ and *Nes*^Cre^hy*TAF1*^XDP^ female mice to orient downward on the pole. ns, non-significant, linear mixed-effects model with age and timepoint as within-subject factors and genotype as a between-subject fixed effect. **i**. Average turn time (s) for male *Nes*^Cre^ and *Nes*^Cre^hy*TAF1*^XDP^ mice in the pole test at week 3. p=0.016, unpaired t-test. **j**. Average turn time (s) for female *Nes*^Cre^ and *Nes*^Cre^hy*TAF1*^XDP^ mice in the pole test at week 3. ns, non-significant, unpaired t-test. **k**. Average force (g) exerted by fore paws by *Nes*^Cre^ and *Nes*^Cre^hy*TAF1*^XDP^ male mice. p < 0.001, linear mixed-effects model with age and timepoint as within-subject factors and genotype as a between-subject fixed effect. **l**. Average force (g) exerted by fore paws by *Nes*^Cre^ and *Nes*^Cre^hy*TAF1*^XDP^ female mice. ns, non-significant, linear mixed-effects model with age and timepoint as within-subject factors and genotype as a between-subject fixed effect. **m**. Average force (g) exerted by fore paws and all paws (fore+hind) by *Nes*^Cre^ and *Nes*^Cre^hy*TAF1*^XDP^ male mice at week 3. Fore paws: p=0.004, All paws, ns=non-significant, unpaired t-test. **n**. Average force (g) exerted by fore paws and all paws (fore+hind) by *Nes*^Cre^ and *Nes*^Cre^hy*TAF1*^XDP^ female mice at week 3. Fore paws: p=0.045, All paws: ns=non-significant, unpaired t-test.

**Supplemental Figure S4.**
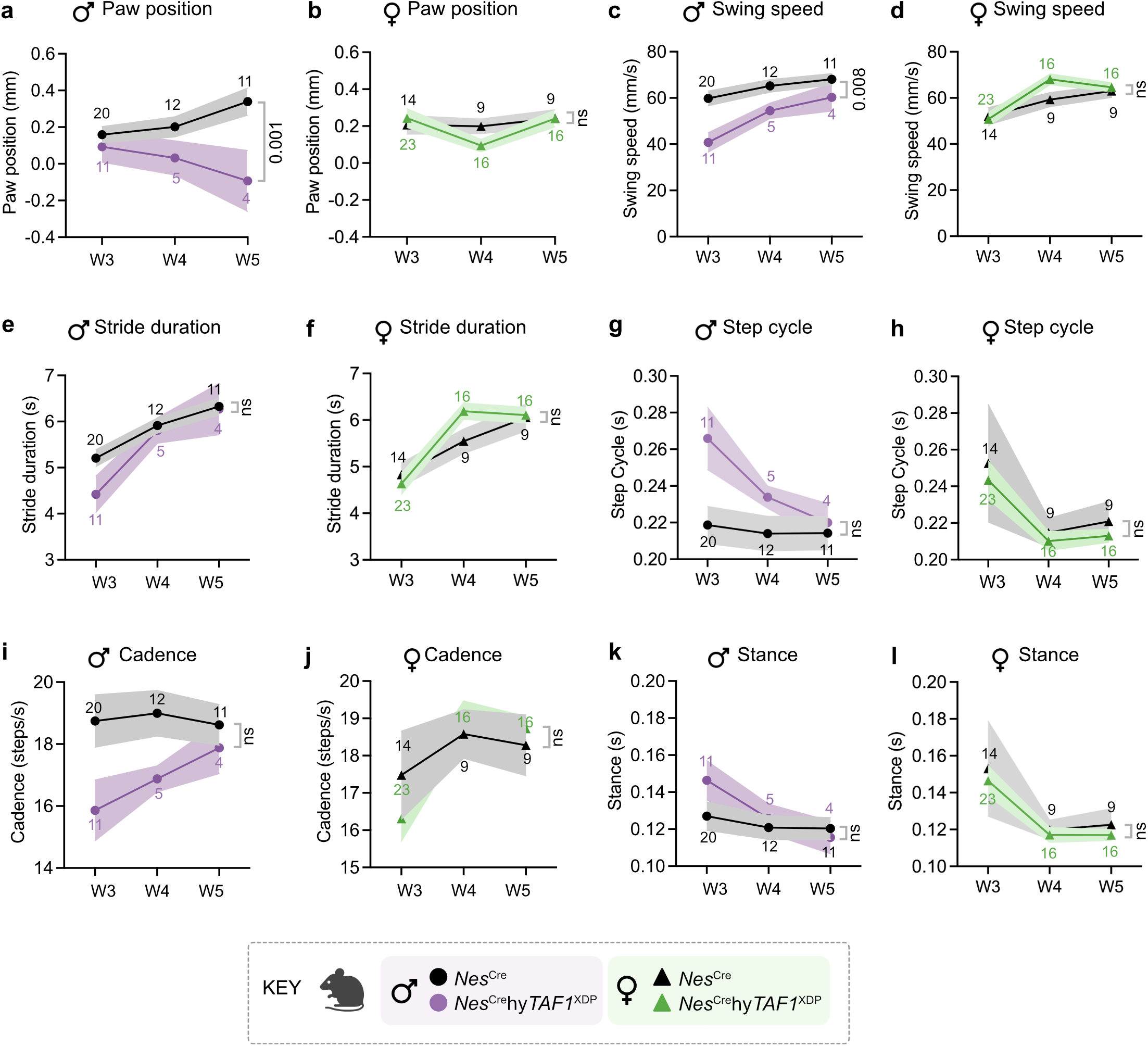
CatWalk™ gait analysis of *Nes*^Cre^hy*TAF1*^XDP^ male and female mice. **a**. Paw position (mm) in male mice. Genotype had a significant effect (F(1, 16.93) = 14.86, p = 0.001), with *Nes*^Cre^ control mice showing higher paw position than *Nes*^Cre^hy*TAF1*^XDP^ mice. Timepoint (TP) and the genotype × TP interaction were not significant (TP: F(2, 14.86) = 0.93, p = 0.415; Genotype × TP: F(2, 14.56) = 2.55, p = 0.113). **b**. Paw position (mm) in female mice. Paw position changed significantly over time (F(2, 25.67) = 6.27, p = 0.006). However, genotype and the genotype × time point interaction were not significant (Genotype: F(1, 23.31) = 0.01, p = 0.917; Genotype × TP: F(2, 26.00) = 0.32, p = 0.732). **c**. Swing speed (mm/s) in male mice (distance covered during swing phase/swing time). Genotype had a significant effect on swing speed (F(1, 16.08) = 9.06, p = 0.008), with *Nes*^Cre^ controls showing higher swing speed than *Nes*^Cre^hy*TAF1*^XDP^ mice. Timepoint (F(2, 12.24) = 1.23, p = 0.327) and the genotype × timepoint interaction (F(2, 12.38) = 0.75, p = 0.493) were not significant. **d**. Swing speed (mm/s) in female mice (distance covered during swing phase/swing time). Effects of genotype (F(1, 25.58) = 0.04, p = 0.849), timepoint (F(2, 24.39) = 0.97, p = 0.393), or genotype × timepoint interaction (F(2, 23.97) = 1.17, p = 0.327) were not significant. **e**. Stride duration (s) in male mice (stand time+swing time). Stride duration changed significantly over time (F(2, 14.37) = 20.26, p < 0.001), increasing between time points W3, W4, and W5. Genotype (F(1, 21.98) = 0.36, p = 0.557) and the genotype × time point interaction (F(2, 13.92) = 1.13, p = 0.351) were not significant. **f**. Stride duration (s) in female mice (stand time+swing time). Stride duration changed significantly over time (F(2, 25.62) = 21.51, p < 0.001), increasing between time points W3, W4, and W5. Genotype (F(1, 29.48) = 0.01, p = 0.907) and the genotype × time point interaction (F(2, 23.89) = 1.81, p = 0.185) were not significant. **g**. Step cycle duration (s) in male mice (stand phase (stance)+swing phase). Step cycle duration changed significantly over time (F(2, 14.53) = 5.99, p = 0.013), increasing across time points W3, W4, and W5. However, genotype (F(1, 22.45) = 0.58, p = 0.455) and the genotype × time point interaction (F(2, 14.03) = 0.03, p = 0.972) were not significant. **h**. Step cycle (s) in female mice (stand phase (stance)+swing phase). There was no significant main effect of time point (F(2, 11.34) = 0.82, p = 0.460), genotype (F(1, 19.11) = 0.41, p = 0.530), or genotype × time point interaction (F(2, 10.18) = 2.62, p = 0.121). **i**. Cadence (steps/s) in male mice (no. of steps/time). Cadence changed significantly over time (F(2, 17.95) = 14.64, p < 0.001), decreasing between time points W3, W4, and W5. Genotype (F(1, 27.12) = 0.26, p = 0.612) and the genotype (× time point interaction (F(2, 15.10) = 0.76, p = 0.485) were not significant. **j**. Cadence (steps/s) in female mice (no. of steps/time). Cadence changed significantly over time (F(2, 25.82) = 11.86, p < 0.001), with pairwise comparisons showing decreases between time points W3 and W4 (p < 0.001) and time points W3 and W5 (p < 0.001). Genotype (F(1, 34.33) = 0.17, p = 0.679) was not a significant predictor. The genotype × time point interaction showed a trend toward significance (F(2, 24.71) = 3.24, p = 0.056). **k**. Stance duration (ms) (time from paw contact to paw lift-off) in male mice. Time point also had a significant effect (F(2, 13.03) = 7.81, p = 0.006). Genotype and the genotype × time point interaction were not significant (Genotype: F(1, 24.07) = 1.07, p = 0.311; Genotype × TP: F(2, 12.33) = 0.09, p = 0.913). **l**. Stance duration (ms) (time from paw contact to paw lift-off) in female mice. Genotype and time point were not significant (Genotype: F(1, 16.48) = 0.53, p = 0.477; TP: F(2, 8.37) = 0.04, p = 0.962). The genotype × time point interaction trended toward significance but did not reach statistical threshold (F(2, 7.36) = 3.24, p = 0.098). Each behavioral measure was analyzed in male or female mice across three timepoints using a linear mixed effects model with genotype, timepoint, and their interaction as fixed effects, and average speed as a covariate. Timepoint (TP) was modeled as a repeated measure with an unstructured covariance matrix. Statistics were derived from a linear mixed effects model with fixed effects tested using Type III F-tests. Plots display significance (p) values or “ns” for genotype comparisons. Number of mice at each time point are displayed on each plot.

**Supplemental Figure S5.**
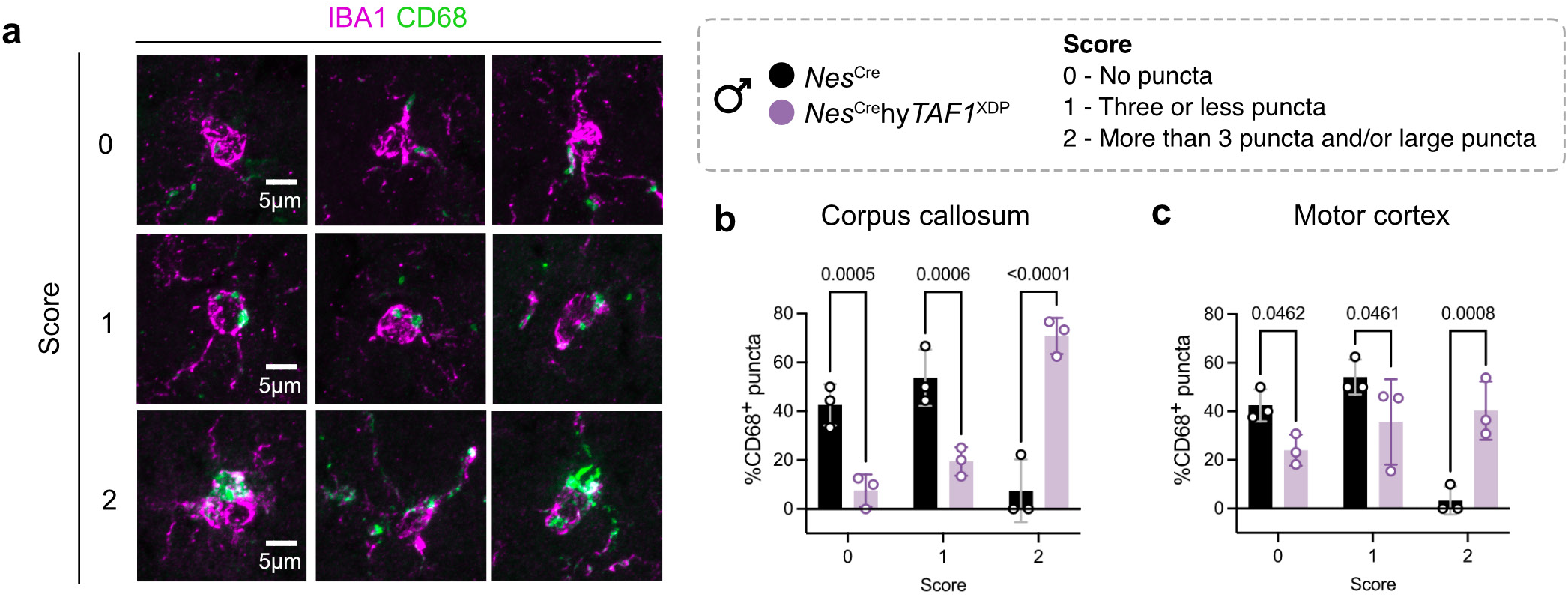
Quantitative analysis of CD68+ puncta in IBA1+ microglia across brain regions. **a**. Representative images showing CD68 puncta scoring in IBA1+ microglia. Microglia were scored based on the abundance and size of CD68+ puncta: Score 0 = no detectable puncta per cell, score 1 = 1-3 puncta per cell, score 2 = more than 2 puncta and/or enlarged puncta per cell. Images illustrate each scoring category. Scale bars = 10 µm. **b**. CD68 puncta scores in the corpus callosum. Mean puncta scores were calculated per mouse across multiple microglia in the corpus callosum. Data are presented as mean ± SEM for n = 3 mice per genotype. p values are shown on the plot, ns: non-significant, repeated measures two-way ANOVA with Šídák’s multiple comparisons test. **c**. CD68 puncta scores in the motor cortex. Mean puncta scores were calculated per mouse across multiple microglia in the motor cortex. Data are presented as mean ± SEM for n = 3 mice per genotype. p values are shown on the plot, ns: non-significant, repeated measures two-way ANOVA with Šídák’s multiple comparisons test.

**Supplemental Figure S6.**
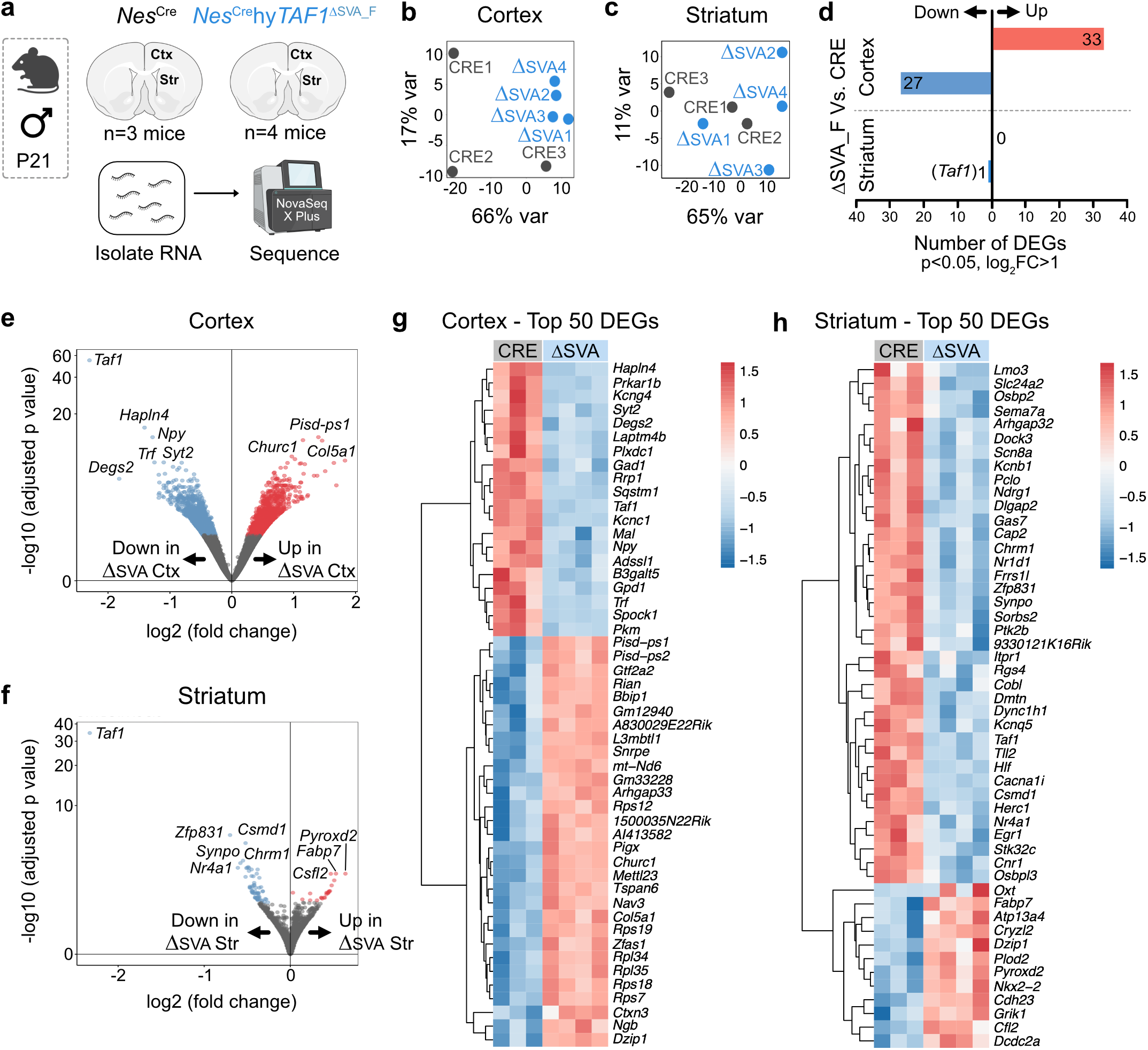
Transcriptomic analysis of *Nes*^Cre^ and *Nes*^Cre^hy*TAF1*^ΔSVA_F^ cortical and striatal tissue by bulk RNA sequencing. **a**. Schematic outlining the experimental workflow for bulk RNA sequencing. Cortical and striatal tissue was micro-dissected from P21 male *Nes*^Cre^ (n=3 mice) and *Nes*^Cre^hy*TAF1*^ΔSVA_F^ mice (n=4 mice). Total RNA was extracted and subjected to bulk RNA sequencing. **b**. Principal component analysis (PCA) of cortical transcriptomes shows no separation between *Nes*^Cre^ (CRE) and *Nes*^Cre^hy*TAF1*^ΔSVA_F^ (ΔSVA) samples, indicating no transcriptional differences between the two genotypes. **c**. Principal component analysis (PCA) of striatal transcriptomes shows no separation between *Nes*^Cre^ (CRE) and *Nes*^Cre-^ hy*TAF1*^ΔSVA_F^ (ΔSVA) samples, indicating no transcriptional differences between the two genotypes. **d**. Bar plot showing the number of differentially expressed genes (DEGs) in the cortex and striatum (adjusted p < 0.05, log_2_ fold change > 1). Red bars represent upregulated genes; blue bars represent downregulated genes. There were no upregulated genes and only 1 downregulated gene (*Taf1*) in the striatum. **e**. Violin plot showing the distribution of log_2_ fold changes and −log_10_ adjusted p values for DEGs in the cortex. Selected genes (*Taf1, Syt12*, etc.) are highlighted. **f**. Violin plot showing the distribution of log_2_ fold changes and −log_10_ adjusted p values for DEGs in the striatum, with select genes labeled. **g**. Heatmap showing the top 50 DEGs in the cortex ranked by adjusted p-value. Expression values are z-scored per gene across all samples. **h**. Heatmap of the top 50 DEGs in the striatum ranked by adjusted p-value. Expression values are z-scored per gene across all samples.

**Supplemental Figure S7.**
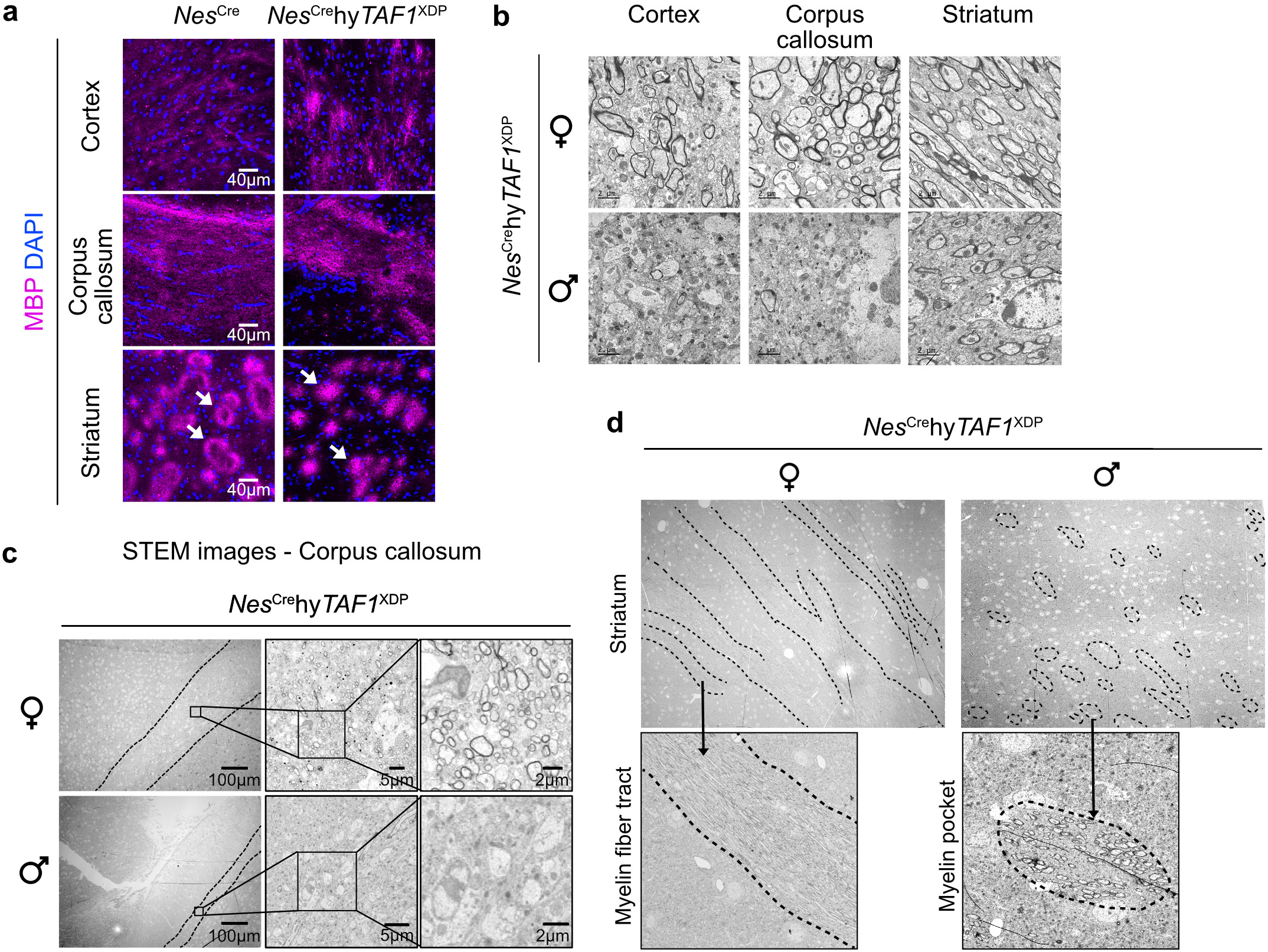
Myelin pathology in XDP mouse brains revealed by immunohistochemistry and electron microscopy. **a**. Immunofluorescence staining for Myelin Basic Protein (MBP, magenta) in the motor cortex, corpus callosum, and striatum of P21 *Nes*^Cre^ and *Nes*^Cre^hy*TAF1*^XDP^ mice reveals disrupted myelin organization in XDP brains compared to controls. Scale bars = 40 µm. **b**. Scanning transmission electron microscopy (STEM) of the corpus callosum from female (top) and male (bottom) *Nes*^Cre^hy*TAF1*^XDP^ mice. Regions of interest with high myelin density are outlined in red. Male *Nes*^Cre^hy*TAF1*^XDP^ brains exhibit a marked reduction in myelin intensity. Scale bars: whole field = 100 µm; inset 1 = 5 µm; inset 2 = 2 µm. **c**. STEM imaging of the striatum in female (top) and male (bottom) *Nes*^Cre^hy*TAF1*^XDP^ brains. Female brains display normal striated patterns typical of myelinated fiber tracts, whereas male brains lack this organization and instead contain discrete, irregular pockets of myelin. **d**. Transmission electron microscopy (TEM) images showing myelin ultrastructure in the cortex, corpus callosum, and striatum from female (top) and male (bottom) *Nes*^Cre^hy*TAF1*^XDP^ mice. Male brains display notable disruptions in myelin sheath integrity. Scale bars = 2 µm.

**Supplemental Figure S8.**
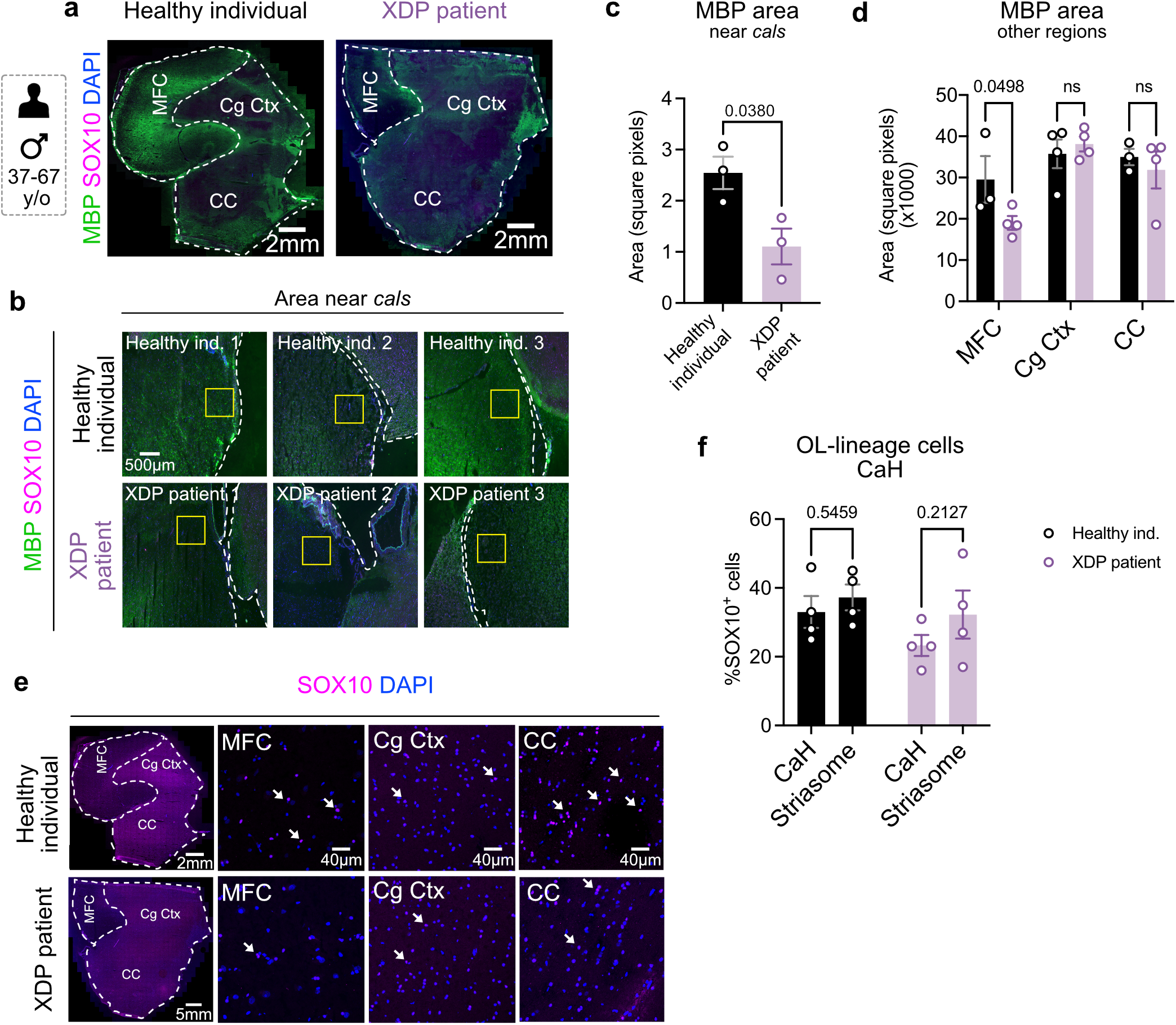
Myelin pathology in postmortem brain tissue from XDP patients. **a**. Immunostaining of medial prefrontal cortex (MFC) near the callosal sulcus (cals) in postmortem brain sections from 43–63-year-old male XDP patients and age-matched healthy controls, showing Myelin Basic Protein (MBP, green) and the oligodendrocyte lineage marker SOX10 (magenta). Scale bars: whole section = 2 mm. **b**. Immunostaining for SOX10+ oligodendrocyte lineage cells in additional brain regions: MFC (away from cals), cingulate cortex (Cg Ctx), and corpus callosum (CC), from XDP patients. White arrows indicate SOX10+ cells. Scale bar = 100 µm. **c**. Overview images showing the region of the MFC near the cals (yellow box) used for myelin analysis. This region shows near-complete loss of MBP staining in XDP patient tissue. Scale bar = 500 µm. **d**. Quantification of MBP+ area in the MFC near cals. XDP patients exhibit a significant reduction in myelin compared to healthy controls. Unpaired t-test; p = 0.0211. Data shown as mean ± SEM; n = 3 individuals per group. **e**. Quantification of MBP area in additional brain regions (MFC away from cals, Cg Ctx, and CC). Two-way ANOVA with Sidak’s multiple comparisons test, ns: nonsignificant. Data shown as mean ± SEM; n = 3 individuals per group. **f**. Quantification of %SOX10+ oligodendrocyte lineage cells in the head of the caudate (CaH) region. Ns, non-significant, two-way ANOVA. Data shown as mean ± SEM, n = 4 individuals per group.

